# Cell growth rates coordinate across the width of the leaf to remain flat

**DOI:** 10.1101/2022.11.01.514736

**Authors:** Kate Harline, Brendan Lane, Antoine Fruleux, Gabriella Mosca, Sören Strauss, Nik Tavakolian, James W. Satterlee, Chun-Biu Li, Abhyudai Singh, Arezki Boudaoud, Richard S. Smith, Adrienne H.K. Roeder

## Abstract

The growth and division of cells in plant leaves is highly dynamic in time and space, even though cells cannot move relative to their neighbors. Thus, organ shape must emerge from carefully coordinated growth, especially in leaves that remain relatively flat as they grow. Here we explored the phenotype of the *jagged and wavy* (*jaw-D*) mutant in *Arabidopsis thaliana*, in which the leaves do not remain flat. It has previously been shown that the *jaw-D* mutant phenotype is caused by the overexpression of *miR319*, which represses TCP transcription factors, thus delaying maturation of the leaf. We analyzed cell dynamics in wild type and *jaw-D* by performing time lapse live imaging of developing leaves. We found that the progression of maturation from the tip of the leaf downward was delayed in *jaw-D* relative to wild type based on several markers of maturation, in agreement with the role of TCP transcription factors in promoting maturation. We further found that these changes in maturation were accompanied by differences in the coordination of growth across the leaf, particularly across the medial-lateral axis, causing growth conflicts that prevent the leaf from remaining flat. Although leaf flatness is often framed as a problem that requires the local synchronization of growth on the abaxial vs adaxial sides of the leaf, our results based on the *jaw-D* phenotype suggest that wild-type plants also need to coordinate growth more globally across the leaf blade to maintain flatness.

## Introduction

In nature, leaves exhibit a myriad of shapes and sizes. Yet, all leaves initially begin as a few cells recruited from the shoot apical meristem (SAM) [1–5]. Understanding how these few initial cells create recognizable forms in nature requires a combined exploration of growth dynamics, tissue mechanics, and genetic regulatory pathways. Imaging at multiple scales has revealed that development and growth in leaves is highly spatially and temporally dynamic [6–11]. In many species, leaves exhibit a dynamic progression from fast to slow growth with a gradient of slow growth initiating from the tip and progressing downward, called the basipetal gradient. The slowing of growth from the tip corresponds with a wave of cellular differentiation that initiates at the tip [7,11–13].

Biomechanical principles predict that juxtaposition of different growth rates within a tissue can create growth conflicts, resulting in deformation of the tissue, including leaf curling and ruffling [14–17]. There are two major types of growth conflict in growth rate and/or growth direction: (1) growth conflict within a tissue layer, and (2) growth conflicts between two cell layers of a tissue, such as adaxial (top) and abaxial (bottom) sides. Neglecting directional growth conflicts, the other two cases occur when cells within a tissue grow at a higher rate than neighboring cells and the region with enhanced growth is compressed by the neighborhood [18]. The compressive stress causes the tissue to buckle and either bend upward or downward. Loss of coordination of growth between the adaxial and abaxial sides of the sepal can cause the tissue to curve or in more extreme cases to make multiple complex ridges through buckling [19,20]. Buckling induced by internal growth conflicts has been used to explain three-dimensional form of fronds in kelp [21], petals in the snapdragon [22], or wing imaginal discs in the fruit fly [23]. Yet, leaves are usually flat, which is optimal for their physiological functions of photosynthesis and gas exchange [1]. It has been shown that a thin elastic sheet growing in three-dimensional space undergoes buckling and loses flatness if heterogeneity in growth is larger than a threshold, which may lead to fractal-like three-dimensional shapes [16,24–27], as observed for leaves of ornamental cabbage. Thus, it is not well understood how leaves maintain their flat form during development.

The *jagged and wavy* (*jaw-D*) mutant is one of the classic *Arabidopsis thaliana* mutants that disrupt leaf flatness [28,29]. *jaw-D* exhibits curly and wavy leaves with ruffles along the edges. The *jaw-D* wavy leaf phenotype has previously been attributed to extended cell division from an over-active marginal meristem [7,30–33]. *jaw-D* was discovered in an activation tagging screen and the phenotype is caused by overexpression of the *microRNA319* (*miR319*) [28,29]. Over the last 20 years, the molecular mechanism through which miR319 regulates its targets has been worked out in exquisite detail; however, it is still not well understood how this pathway contributes to leaf flatness. miR319 targets the class II *TEOSINTE BRANCHED 1, CYCLOIDEA, PROLIFERATING CELL NUCLEAR ANTIGEN FACTOR 1 AND 2 (TCP)* transcription factors which have been shown to repress growth and activate tissue differentiation [31,32,34,35]. miR319 targets, especially *TCP4,* are thought to regulate cell growth and differentiation through *CUP-SHAPED COTYLEDONS 2 (CUC2)* and the *CUC* regulator miR164 as well as auxin and cytokinin networks [36–40]. Based on GUS assays, previous studies have shown that the promoters of genes in the miR319 family drive expression at the base of leaves (miR319c) and floral organs (miR319a,c) [39,41]. Thus, the evidence suggests that miR319 and TCPs contribute to the wave of differentiation and cessation of cell division from the tip of the leaf downward. Still, it is unclear how this proximal-distal wave of differentiation relates to maintenance of leaf flatness.

The regulation of TCPs by miR319 is well-conserved within angiosperms [42,43]. Therefore, this pathway could have broad implications for the flattening of leaves throughout the angiosperms. Notable homologs in other model species include the class II-defining member *CINCINNATA (CIN)* in snapdragon and *Lanceolate* in tomato [44–46], both of which when mutated or downregulated yield rippling and undulating leaves like the Arabidopsis *miR319* overexpression mutant *jaw-D.* Increased curvature in *cin* leaves was attributed to decreased perception of a tip-derived differentiation signal and thus an escape from leaf margin growth regulation [44] . Still, the connection between organ shape and a proximal-distal differentiation wave regulating cell growth remains to be resolved.

To understand how leaf flatness emerges, we have tracked how the cell growth, cell division and cell maturation give rise to the bottom (abaxial) epidermis of the developing leaf blade in living wild-type and *jaw-D* mutant first leaves. Because *jaw-D* overexpresses miR319, we expected to see a disruption of the proximal-distal growth gradient. We confirmed this disruption, finding that growth and cell division were spread more distally in *jaw-D,* but with a lower maximum value than wild type. Although we did not observe overgrowth of the leaf margins in *jaw-D* relative to wild type in the first leaves, which was expected based on previous work, we did observe that cell growth across the medial-lateral axis of the leaf was less coordinated and more patchy than in wild type. In particular, growth was coordinated between the midrib and leaf blade in wild type, but less so in *jaw-D*. Modeling revealed that coordination of growth along the medial-lateral axis, while still maintaining the growth gradient from tip to base was sufficient to create a flat leaf blade, similar to wild type. In the model, lack of coordination of growth between midrib and blade created a curved leaf similar to *jaw-D*. Thus, our results suggest wild-type plants coordinate growth across the medial-lateral axis leaf blade to maintain flatness.

## Results

### The generation of leaf shape can be captured through live imaging early stages of leaf development

To understand how the three-dimensional form of the leaf arises from cellular growth patterns, we repeatedly imaged the same living wild-type and *jaw-D* mutant first leaves every day from 3 days after sowing (DAS) to 8 DAS (n = 3 leaves for each genotype; Fig 1A; Supplemental Figs S1-S6). The first two leaves develop simultaneously and are indistinguishable, so we consider them together as leaf 1 and 2. To increase the surface of the leaf that could be imaged, the cotyledons were dissected off [47] . We used confocal microscopy to image nearly the entire abaxial (bottom) epidermis of the leaf at cellular resolution so that we could relate cellular properties to organ scale measurements. Then we used MorphoGraphX 2.0 image analysis software to represent the confocal stacks as two dimensional mesh surfaces curved in three dimensions (2.5D) [48,49]. Cell boundaries were marked with a fluorescent plasma membrane marker. We segmented the epidermal cell signal projected on this mesh and tracked each of the cell lineages over time (Video S1-S4). This dataset serves as the basis for our further analysis of how cell growth and division give rise to organ shape (Dataset available at https://doi.org/10.17605/OSF.IO/D7X3Y).

**Fig 1.**
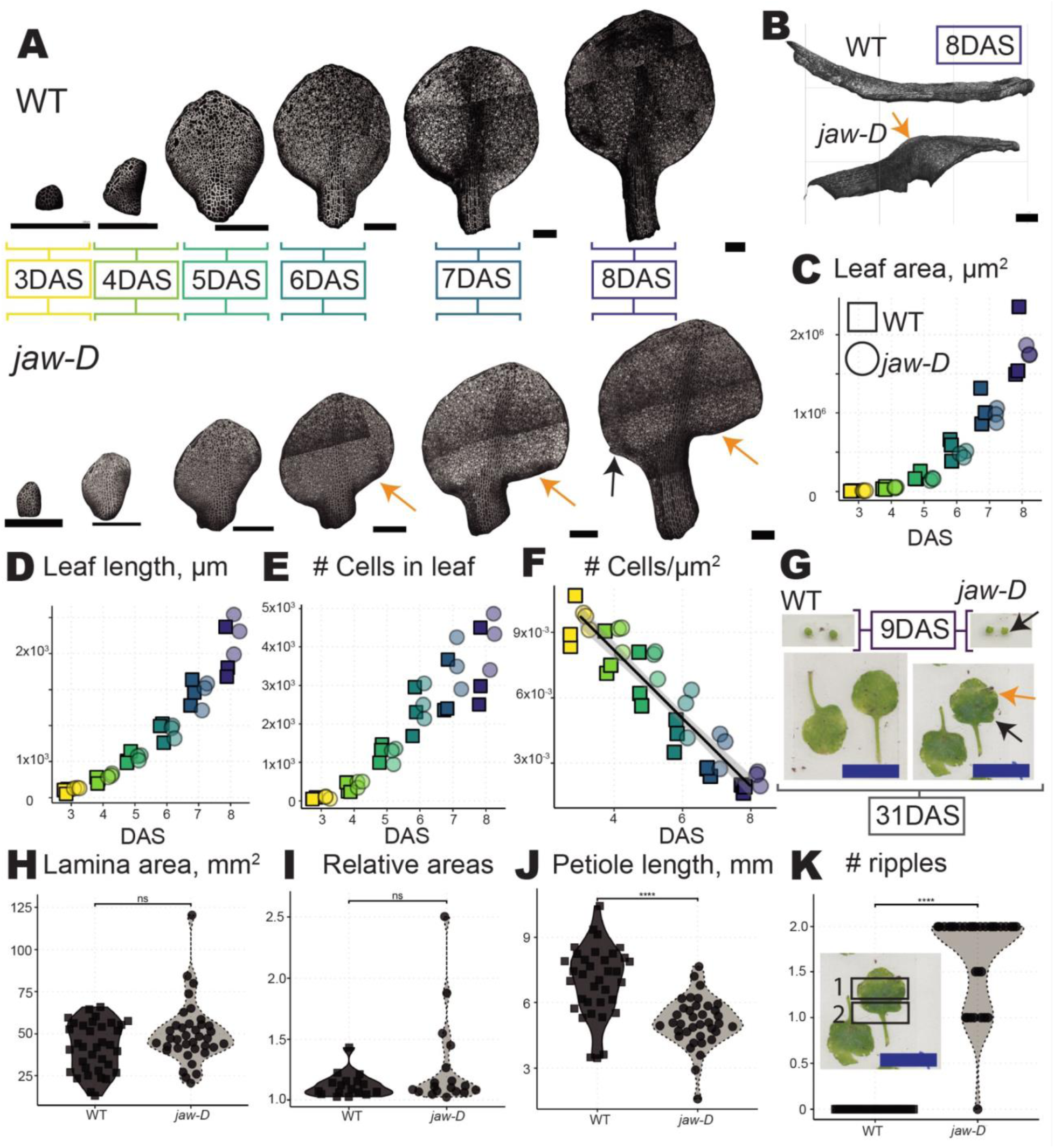
Features of *jaw-D* leaf form emerge soon after primordia initiation. (A) Fluorescent plasma membrane marker signal from confocal stack images of wild type (WT; top) and *jaw-D* (bottom) leaves from 3 DAS to 8 DAS are shown projected onto 2.5D surface meshes rendered in MorphoGraphX 2.0. Tissue differentiation in WT and its disruption in *jaw-D* can be seen starting at 5 DAS. Black arrows indicate the beginnings of leaf dimples and orange arrows indicate increased curvature in early *jaw-D* leaves that will become ripples at maturity. (B) Side view of 8 DAS leaf meshes, with petiole on the left and tip on the right, show a global increase in leaf curvature in *jaw-D* (lower) versus WT (upper). (C-D) Total segmented leaf area (C) and length (D) increase exponentially over the course of the live imaging experiment in WT and *jaw-D* plants. (E) The number of segmented cells in WT and *jaw-D* begins to plateau around 6 DAS, suggesting that the major phase of cell division is ending. (F) Cell density was determined as the number of cells per area segmented. The density is mostly linear from 3-7 DAS in *jaw-D* and WT but begins to show asymptotic behavior reflective of plateauing cell numbers. Squares indicate WT leaves, circles indicate *jaw-D* leaves. Colors indicate DAS. Means for all measures for each time point were not significantly different (paired Student’s t-tests with Bonferroni correction). n=3 leaves for each genotype. Black scale bars = 200 µm. (G) Comparison of the first and second leaves of WT (left) and *jaw-D* (right) from 9 DAS (top) and mature leaves, at least 31 DAS (bottom). Blue scale bars indicate 10 mm. (H) Final lamina area for the first and second leaves of WT and *jaw-D*. WT leaf areas exhibit a broad and bimodal distribution, whereas *jaw-D* leaf areas center tightly around the median with more extreme values (p > 0.05, Wilcox test). (I) Relative area between the final sizes of the first and second leaves of the same plant in WT or *jaw-D. jaw-D* first and second leaves on the same plant have a greater spread of values and more extreme values (Feltz and Miller asymptotic test of CV p = 3.051466e-05). (J) Length of petioles in fully grown first and second leaves of WT and *jaw-D. jaw-D* petioles are shorter (p < .0001, Wilcox test). (K) Number of ripples in mature *jaw-D* plants (versus none in WT p < .0001, Wilcox test). Inset indicates identification of ripples in mature *jaw-D* plants. Color, shape and linetype indicate genotype (solid, dark gray, squares = WT, dotted, light gray, circles = *jaw-D*). See supplemental Figs S1-S6 for three replicate wild type and *jaw-D* leaves showing the change in size, shape, and differentiation from 3 DAS to 8 DAS. See Supplemental Fig S7 for further quantification of wild-type and *jaw-D* leaf growth.

To ensure that we had captured the relevant growth events that establish leaf flatness in wild type and curvature in *jaw-D*, we examined large-scale leaf development in our images during the 3-8 DAS that we imaged. Starting at 6 DAS tissue curving became evident in *jaw-D* leaves, while wild-type leaves remained relatively flat (Fig 1A-B). This tissue curving continued throughout the experiment and was accompanied by ruffling along the margin at 8 DAS (Fig 1A-B, G, Video S5). In all samples, we observed rapid cellular and organ-wide morphology changes characteristic of early leaf growth (Fig 1A, Supplemental Figs S1-S6). Notably, at 4 DAS stomata began to initiate at the distal end of the leaf in both wild type and *jaw-D*, indicating that the basipetal (distal to proximal) maturation and differentiation gradient had initiated at the distal leaf tip. At 5 DAS the petiole (stalk that attaches the leaf to the stem) initiated as a cylindrical fold in the proximal base of the leaf, differentiating it from the lamina (the flat leaf surface) in both wild-type and *jaw-D* leaves. Thus, in the first 8 days of leaf development, we have captured key changes in cellular growth that cause the formation of curvature in the *jaw-D* leaf shape.

Organ-wide morphological features of the 3-8 DAS leaves, including area, length, width, cell number, and cell density, were not statistically different between wild type and *jaw-D* (paired Student’s t-tests with Bonferroni correction, Fig 1C-F Supplemental Fig S7A). Therefore, we concluded that wild-type and *jaw-D* leaves could be considered at the same stage for each day in the experiment. However, the enhanced curvature of *jaw-D* prevented cell segmentation in some of the proximal leaf blade regions, so it is likely that proximal *jaw-D* cells are undercounted. While leaf area and leaf length continued to increase exponentially throughout the experiment (log(leaf area) ∼ time, R^2^ = 0.93, p < 2.2e-16; log(leaf length) ∼ time, R^2^ = 0.94, p < 2.2e-16, Supplemental Fig S7F-G), cell number began to plateau (three parameter log-logistic Lack of fit p-value = 0.95) around 6 DAS, suggesting that we captured the transition of leaves from growth associated with cell division to growth predominately through cell expansion (Fig 1E) [1] . Additionally, the density, the number of cells per area, stayed mostly linear, but was approaching a negative exponential curve in the last two time points in wild type, suggestive of a more complete transition to expansion-based growth in wild type (cell density ∼ time, R^2^ = 0.92, p < 2.2204e-16; Fig 1F). Thus, with our imaging of 3-8 DAS we captured both the generation of flatness/curvature and the basipetal cell division and growth gradient.

To relate the phenotypes we captured early in leaf development to phenotypes of fully developed leaves, we scanned wild-type and *jaw-D* leaves 1 and 2 at maturity (at least 31 DAS). When we measured the mature first true leaves of wild-type and *jaw-D* plants, *jaw-D* leaves could not be flattened and therefore small tears at the margin had to be induced to measure leaf characteristics, as reported previously (Fig 1G-K) [31]. By measuring paired leaves 1 and 2 from >15 plants each (wild-type n = 34 leaves, *jaw-D* n = 36 leaves), we observed that leaves undergo approximately a 400-fold increase in area from 8 DAS to maturity, yet the gross morphology is established by 8 DAS. Generally, the first two leaves of *jaw-D* form two ripples and these ripples are established during the live imaging experiment window, while wild-type leaves never form ripples (Fig 1A-B, G, K). wild-type and *jaw-D* leaves have the same median size at maturity (Fig 1H, Wilcox test p > 0.05). Other measurements of organ size between wild type and *jaw-D* are likewise comparable (Supplemental Fig S7B-E). We did find that the petiole of *jaw-D* is shorter than wild type (Fig 1J, Wilcox test p < .0001).

To test the robustness of organ size, we next asked whether leaves 1 and 2 are the same size (i.e robust) in a single plant. We divided the area of the larger leaf from the pair by the area of the smaller leaf. If leaves are equal sizes, the result is one. The median value was close to 1 and not statistically significantly different between wild-type and *jaw-D* plants (Fig 1I, 1.09 wild type versus *jaw-D* 1.10, Wilcox test p > 0.05), indicating in most plants the leaves were very close to the same size, and thus robust. However, *jaw-D* exhibited more outlier plants with extremely different leaf 1 and 2 sizes (statistically higher coefficient of variation Feltz and Miller Asymptotic test p = 3.051466e-05), indicative of higher size variability induced by the *jaw-D* mutation. We conclude that jaw-D leaves are more sensitive to perturbations.

We conclude that our live imaging of the early stages of leaf development (3-8 DAS) captures the initial development of organ shape at cellular resolution. We were curious how leaves with approximately the same starting and ending tissue material could have such distinct shapes, namely flattened in wild type and curved in *jaw-D*. In the next sections, we associate the spatial and temporal initiation of this curvature with cellular growth, division and differentiation patterns.

### While wild-type leaves progressively flatten, *jaw-D* leaves curve at the base

To quantify the magnitude and direction of tissue curvature, we calculated the Gaussian curvature for live imaged wild-type and *jaw-D* samples (Fig 2, Supplemental Fig S8). The sign of Gaussian curvature indicates the structure: a hill or valley (positive) or saddle (negative). Values of Gaussian curvature further from zero indicate greater curvature. At 6 DAS, *jaw-D* leaves establish a strong positive curvature across the proximal end of the leaf lamina as the whole leaf curves (Fig 2B, Supplemental Fig S8A). At early time points (3-5 DAS), wild-type leaves exhibit a range of curvature values (Supplemental Fig S8A). At 6 DAS, while *jaw-D* leaves curve, wild-type curvature values along the length and width tighten around 0, as the leaf flattens (Supplemental Fig S8). We conclude that the entire *jaw-D* leaf curves, not just the margins, and that curvature is initiated at 6 DAS.

**Fig 2.**
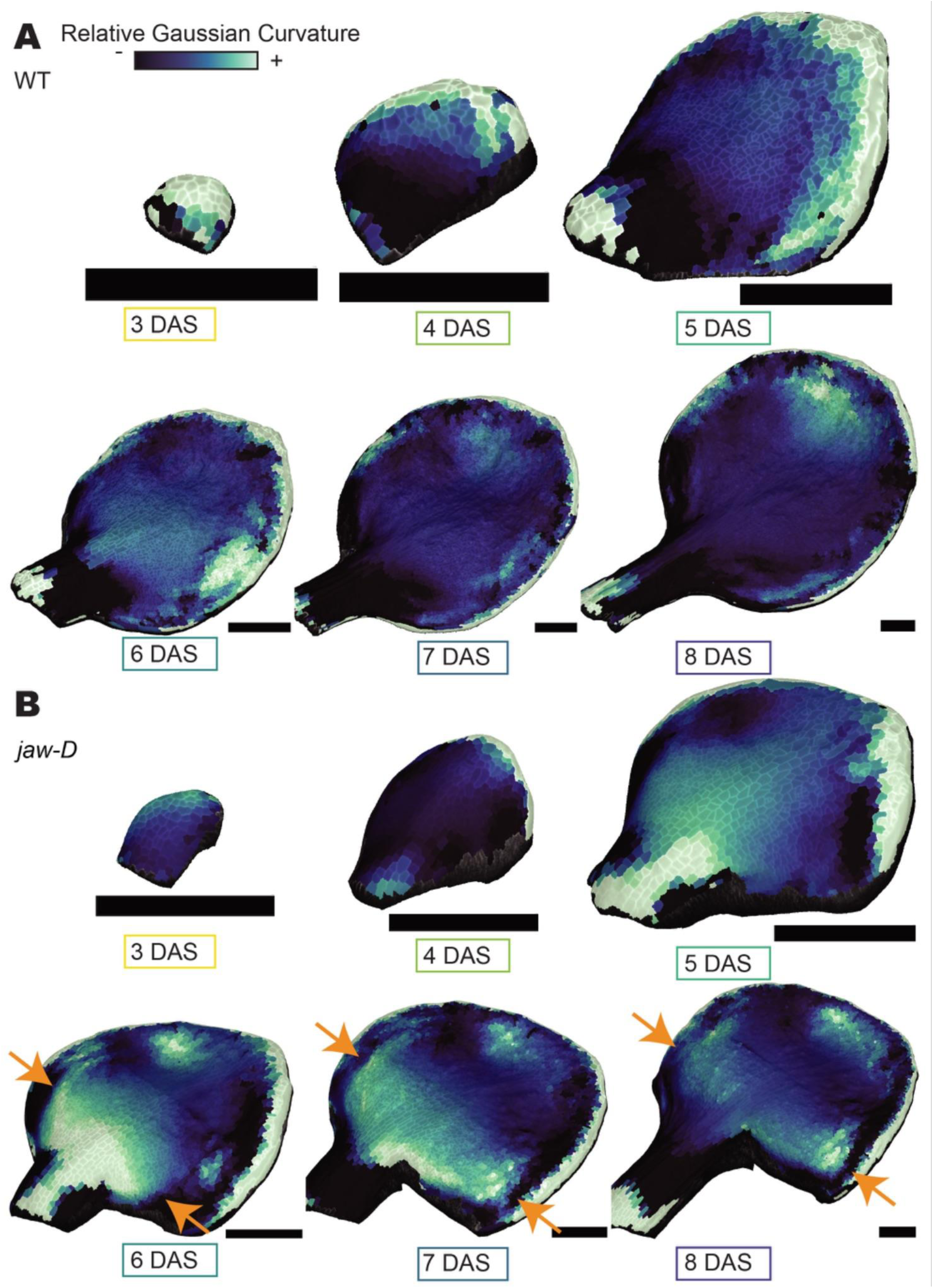
While WT leaves progressively flatten, *jaw-D* leaves curve at the base. (A-B) Gaussian curvature displayed as heatmaps on WT (A) and *jaw-D* (B) leaf meshes. Gaussian curvature represents both the magnitude and direction of curvature such that values further from zero represent greater curvature. Orange arrows indicate high positive curvature that develops at the base of *jaw-D* leaves. Black scale bars = 200 µm. These images are representative of three replicate live imaging series for each WT and *jaw-D*. Note, as all leaves flatten, the Gaussian curvature changes dynamically over multiple orders of magnitude during the experiment. Further, the petiole and margin are sources of extreme outliers in curvature. Therefore, the scale for the heatmaps representing each time point has been calculated to encompass the 85% range of curvature values around the mean for that time point. See Supplemental Figure S8 for quantification of curvature.

### Wild-type leaves exhibit a well-defined proximal-distal gradient of growth and division, which is lacking in *jaw-D*

To investigate the relationship of cells to organ-level curvature, we analyzed the patterns of cellular growth and division in the leaf. From our segmented and lineage tracked cells, we calculated the growth rate (relative change in area) and number of divisions undergone by each cell at each time point in the live imaging experiment (Fig 3A-D). To quantify the relationship between cell division and growth in tandem, we represent each of these values for each cell in its normalized proximal-distal and medial-lateral position (Fig 3E, Supplemental Fig S9).

**Fig 3.**
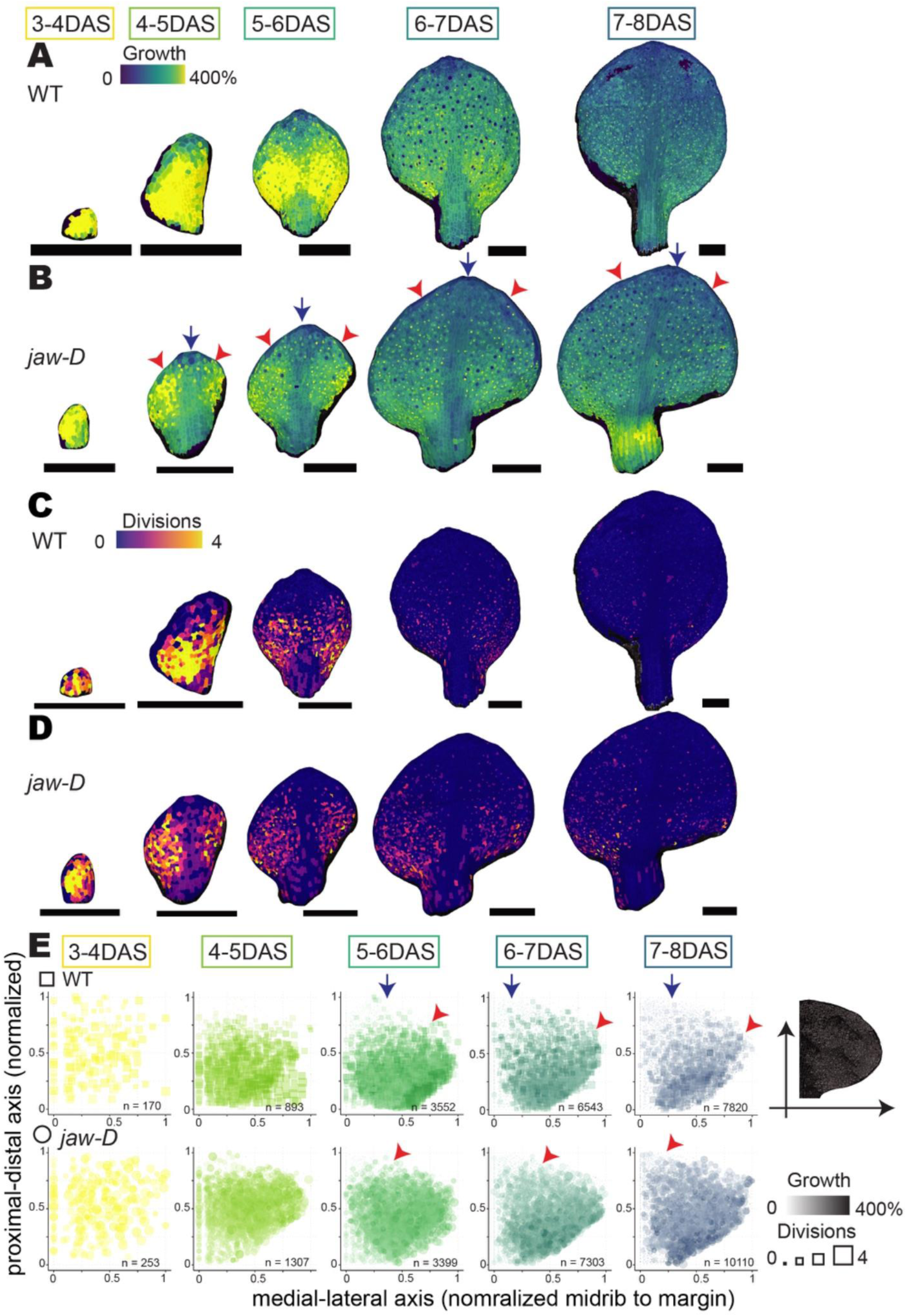
WT leaves exhibit a well-defined basipetal gradient of growth and division and uniform growth from midrib to margin, both lacking in *jaw-D*. (A-B) Heat-maps of cell areal growth (from 400% fast growth in yellow to 0% no growth in navy blue) in 24 hour imaging periods represented on sample WT (A) and *jaw-D* (B) leaf meshes on the earlier time step. Note the slowing of growth that initiates from the distal tip at 4-5 DAS and progresses toward the base in WT. This basipetal gradient is less obvious (somewhat flattened) in *jaw-D*. Note also that in WT the growth rate is the same from midrib to margin at a specific location on the proximal distal axis, whereas in *jaw-D* the midrib consistently grows slower (blue arrows) than the leaf blade (red arrows). (C-D) Heat-maps of the number of cell divisions (from 4 in gold to 0 in purple) in 24 hour imaging periods represented on sample WT (C) and *jaw-D* (D) leaf meshes on the earlier time step. Note the slowing of division that initiates from the distal tip at 4-5 DAS and progresses toward the base in WT. This basipetal gradient is less obvious (somewhat flattened) in *jaw-D*. Black scale bars = 200um. (E) All cells tracked in all three laminas of WT (top, squares) or *jaw-D* (bottom, circles) samples. n indicates the number of cells in each panel. Colors indicate 24 hour time points. The x and y axes represent medial-lateral and proximal-distal positions along the leaf lamina, respectively, normalized between 0 and 1 for the given replicate and time point. The size and color intensity of each point represents the number of divisions and areal growth ratio, respectively, a cell has experienced between imaging days. WT cells show a distinct region of extensive growth and division (dark, big squares) in the lower half of the lamina starting at 5-6 DAS, while *jaw-D* cells exhibit a broader and less intense region in the bottom three quarters of the lamina. Blue arrows indicate the slow growing and dividing region starting from the top of the leaf and expanding downward with the basipetal growth gradient. Red arrows demarcate the boundary between fast and slow growth in the basipetal growth gradient. Note this boundary is more toward the tip of the *jaw-D* leaf and does not progress downward.

Wild-type leaves have previously been shown to have a basipetal (progressing from tip to base) growth gradient, in which slowing growth and cessation of cell division initiate in the tip and progress downward. Thus, at many developmental stages, wild-type leaves exhibit fast growth in the proximal base and slow growth in the distal tip. Given that TCPs promote slowing growth and cessation of cell division to create the basipetal growth gradient in wild-type leaves, we expected the basipetal growth gradient to be disrupted in *jaw-D* because overexpression of the *miR319* inhibits TCPs [31,32,34,35]. As anticipated, the basipetal growth gradient initiates, but does not properly progress down the leaf in *jaw-D* as it does in wild type. In both wild type and *jaw-D* over 3-4 DAS, the whole leaf primordium grows and divides rapidly (Fig 3). At 4-5 DAS in both wild type and *jaw-D*, cells in the distal tip slow their growth and stop dividing, initiating the basipetal growth gradient (Fig 3). In wild type slowing growth and cessation of cell division proceed down the leaf blade over 5-8 DAS (Fig 3A,C,E, Fig 4A-B, Supplemental Fig S9). In contrast, in *jaw-D*, a moderate growth and division rate continues to extend across the leaf blade (Fig 3B,D,E, Supplemental Fig S9, S10A-B). All three leaf replicates for each genotype exhibit these respective growth patterns (Supplemental Fig S9). We conclude that the basipetal growth gradient that is prominent in wild type is perturbed in *jaw-D*.

**Fig 4.**
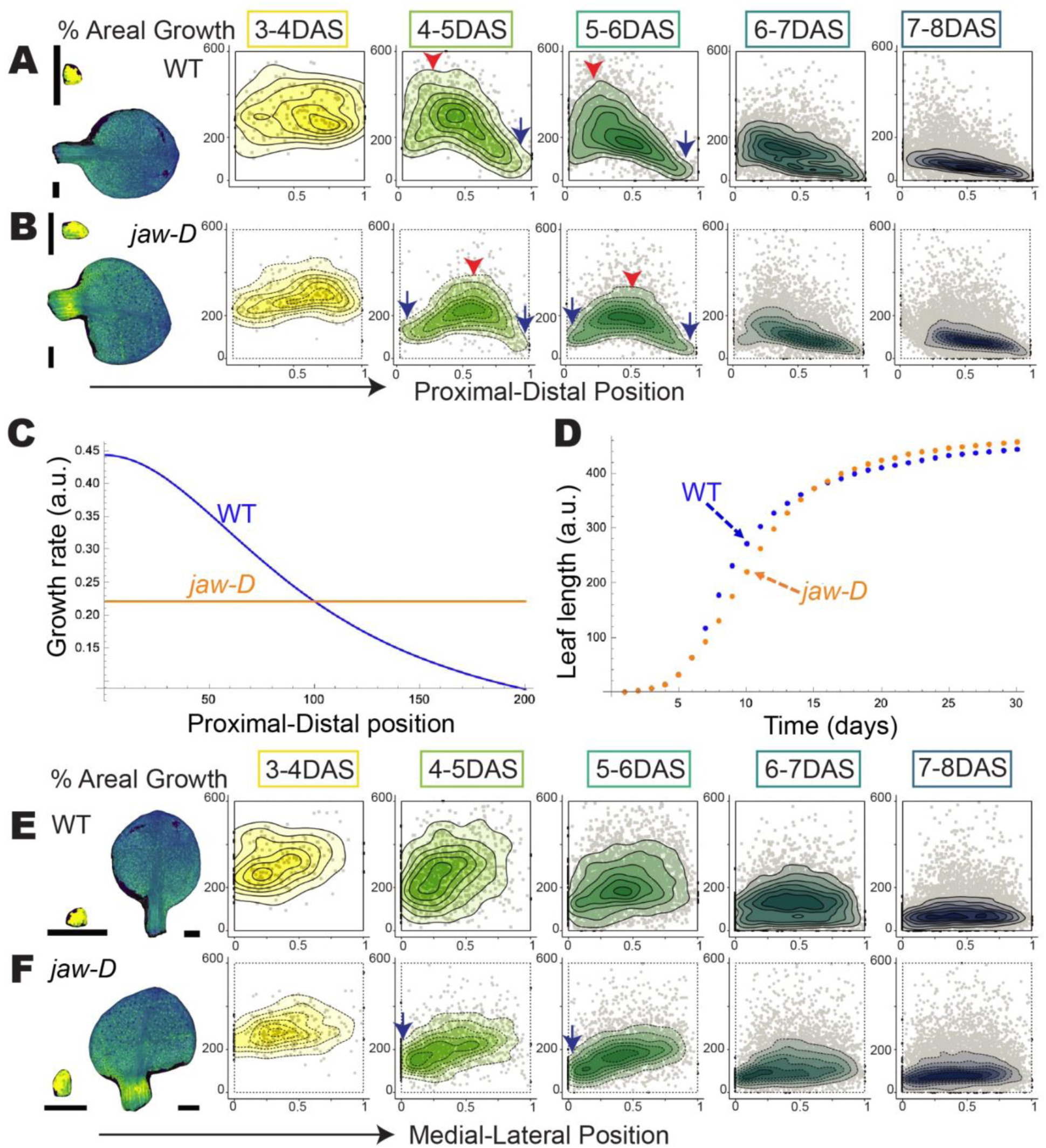
Quantification of disruption in the basipetal growth gradient and midrib to margin growth in jaw-D versus wild type. (A-B) Density plots of each cell’s percent cell area increase in 24 hours graphed according to cell position along the proximal-distal axis (normalized so the tip is 1 and base is 0) to examine the basipetal growth gradient. In day 4-5 and 5-6 there is a pronounced slope with slower growth (blue arrow) in the tip and faster growth in the base (red arrow) of wild type leaves, showing the basipetal growth gradient. At the same time points, the cell area increase in *jaw-D* is more uniform across the length of the leaf with the highest areal growth percentages in the center (red arrow) compared to both ends (blue arrows). The images of the areal growth heat maps (reproduced from Fig. 3) illustrate the alignment of the proximal distal axis in the graphs. Black scale bars = 200 um. (C-D) Leaf length model with growth distribution across the leaf (C) and resultant leaf length (D) represented for WT (blue) and *jaw-D* (orange) simulated values. The growth rate profiles in (C) correspond to a leaf length of 200 a.u. in (D). (E-F) Density plots of each cell’s percent cell area increase in 24 hours graphed according to cell position along the medial-lateral axis (normalized so the margin is 1 and midrib is 0) to examine the uniformity of growth in the medial-lateral direction. Note that from 4-5 and 5-6 DAS the growth near the midrib is lower (blue arrows) than in the blade in *jaw-D*. The images of the areal growth heat maps (reproduced from Fig. 3) illustrate the alignment of the medial-lateral axis in the graphs. Black scale bars = 200 um.

To further quantify the defects in the basipetal growth gradient in *jaw-D*, we examined the level of daily growth and divisions along the proximal-distal axis of the leaf. Wild-type leaves show the classic basipetal growth gradient with low growth at the distal tip and high growth rates near the proximal base from 4-7 DAS (Fig 4A). We found a striking shift in the distribution of areal growth in *jaw-D* leaves along the proximal-distal axis, with growth occurring at a more constant rate all the way along the proximal distal axis. The slight maxima growth in *jaw-D* occurs near the middle of the leaf throughout 4-6 DAS when the change in curvature initiates in *jaw-D* (Fig 4B). Cell divisions showed similar trends (Supplemental Fig S10A-B) with the *jaw-D* distribution centered near the middle half and the wild-type distribution decreasing from a maximum in the proximal quarter of the leaf. Thus, the basipetal growth and division gradient of wild type is flattened in the *jaw-D* leaf. This suppression of the basipetal growth gradient in *jaw-D* is consistent with the current understanding of disruption of the *miR319 TCP* network in *jaw-D*. TCP transcription factors promote maturation from the tip downward [31,32,34,35]. Ectopic expression of *miR319* in *jaw-D* leads to expanded repression of *TCP* expression and consequently is expected to suppress the basipetal growth gradient as observed in our live imaging analysis.

We were struck by the observation that the *jaw-D* leaf does not become longer than wild type (Fig 1D) despite having an expanded region of fast growth and division (Fig 3). To explore how contrasting spatial growth profiles for wild-type and *jaw-D* genotypes can lead to overall similar leaf length dynamics, we created a simple one-dimensional model to capture the length of the leaf over time. In the model, we assume growth is initially exponential in both wild type and *jaw-D*. As the leaf dynamics exit this initial exponential phase, for wild type, we implement a basipetal growth gradient curve (Fig 4C). We also assume that the growth at the leaf base decreases over time; this decrease is essential to eventually stop all growth as the leaf length approaches its steady-state value. For *jaw-D* we simplify the pattern and implement a constant moderate growth rate along the leaf length (Fig 4C). Consistent with our data, the model predicts similar leaf lengths can be maintained between wild type and *jaw-D* despite these different growth profiles (Fig 4D). Based on the model, we conclude that different proximal distal growth patterns can yield similar leaf lengths. Thus, a flattening of the basipetal growth gradient is one of the major defects in *jaw-D* mutants, but it was not yet clear how alterations in *jaw-D* cellular growth affect curvature.

### In *jaw-D* leaves the midrib grows slower than the leaf blade, whereas wild-type growth is more even across the medial-lateral axis

To look for further defects in the cellular growth of *jaw-D* that could give rise to changes in curvature, we also examined growth in the medial-lateral (middle to edge) leaf axis. In previous publications, leaf ruffling has been attributed to overgrowth along the leaf margins [7,30]. We did not observe any obvious overgrowth or excessive division in the edges of *jaw-D* leaves beyond that in wild type (Fig 3). We note that our live imaging captured the first of two ripples that will be formed in the first leaves of *jaw-D* (Fig 1B and G). We also note that we use the term ripple, because the phenotype is not just at the margin, but curvature that extends across the whole leaf. Since curvature is affected across the *jaw-D* leaf, not just the margins, we looked more broadly for defects in growth in the medial-lateral direction. The *jaw-D* midrib (middle of the leaf overlying the central vascular bundle) grows more slowly than the leaf blade from 4-8 DAS (Fig 3B). In contrast, the wild-type midrib and blade have nearly equivalent growth rates along the same latitudinal level (Fig 3A). Further quantifying growth rates along the medial-lateral axis confirmed slower growth near the midrib and faster growth throughout the blade in *jaw-D*, whereas growth rates were more broadly and evenly distributed in wild type (Fig 4E-F). Cell divisions largely cease at 5-6 DAS and are less distinguishable medio-laterally between wild type and *jaw-D* (Supplemental Fig S10C-D). So, we reasoned that growth, rather than divisions, was a major driver in creating the curvature differences in *jaw-D* leaves. Our data is consistent with the hypothesis that differences in growth rate between the midrib and the leaf blade in the medial-lateral axis create mechanical conflicts that cause the rippling of the whole *jaw-D* leaf. In contrast, the relatively even growth rate in the midrib and blade in wild type would not cause conflicts so the leaf can retain flatness.

In summary, our growth analysis revealed two major defects in *jaw-D* growth relative to wild type. First the basipetal growth gradient in the proximal distal direction is flattened in *jaw-D* compared to wild type. Second, in the medial-lateral direction, the midrib growth is slower than the rest of the leaf blade in *jaw-D*, whereas the midrib grows as fast as the blade in wild type. Therefore we examined the growth and development patterns of the leaves along these two axes in more detail to understand how cellular growth gives rise to curvature in *jaw-D* and flatness in wild type.

### Analysis of growth rate variability confirms the two major defects in *jaw-D* growth patterns: flattened basipetal growth gradient and slow growth at the midrib

To further characterize the cellular growth patterns, we quantified the local variability in growth rates, i.e. how different is the growth rate of a cell from its neighbors and how much does the growth rate of an individual cell change between one time point and the next [50]. In both local spatial and temporal variability, *jaw-D* shows a prominent reduction in cell growth variability at the midrib between 5 and 8 DAS (Fig 5A-B). The lower variability in midrib cell growth confirms the persistent slower growth rate of the midrib relative to the more dynamic growth of the leaf blade. In contrast, the midrib is not as distinct in wild type because the midrib cell growth is more equivalent to that in the leaf blade (Fig 5A-B). If we quantify the overall local spatial and temporal variability by averaging cells throughout the leaf, we found that wild type and *jaw-D* are largely indistinguishable (Fig 5C-D), indicating that the localized pattern, not the global pattern is important in the case of medial-lateral growth. Stomata have recently been suggested as the source of most growth heterogeneity in tissues [51], which is also evident in our heat maps (Fig 5A-B). However, our analysis was largely unaffected when stomata were excluded (Supplemental Fig S11).

**Fig 5.**
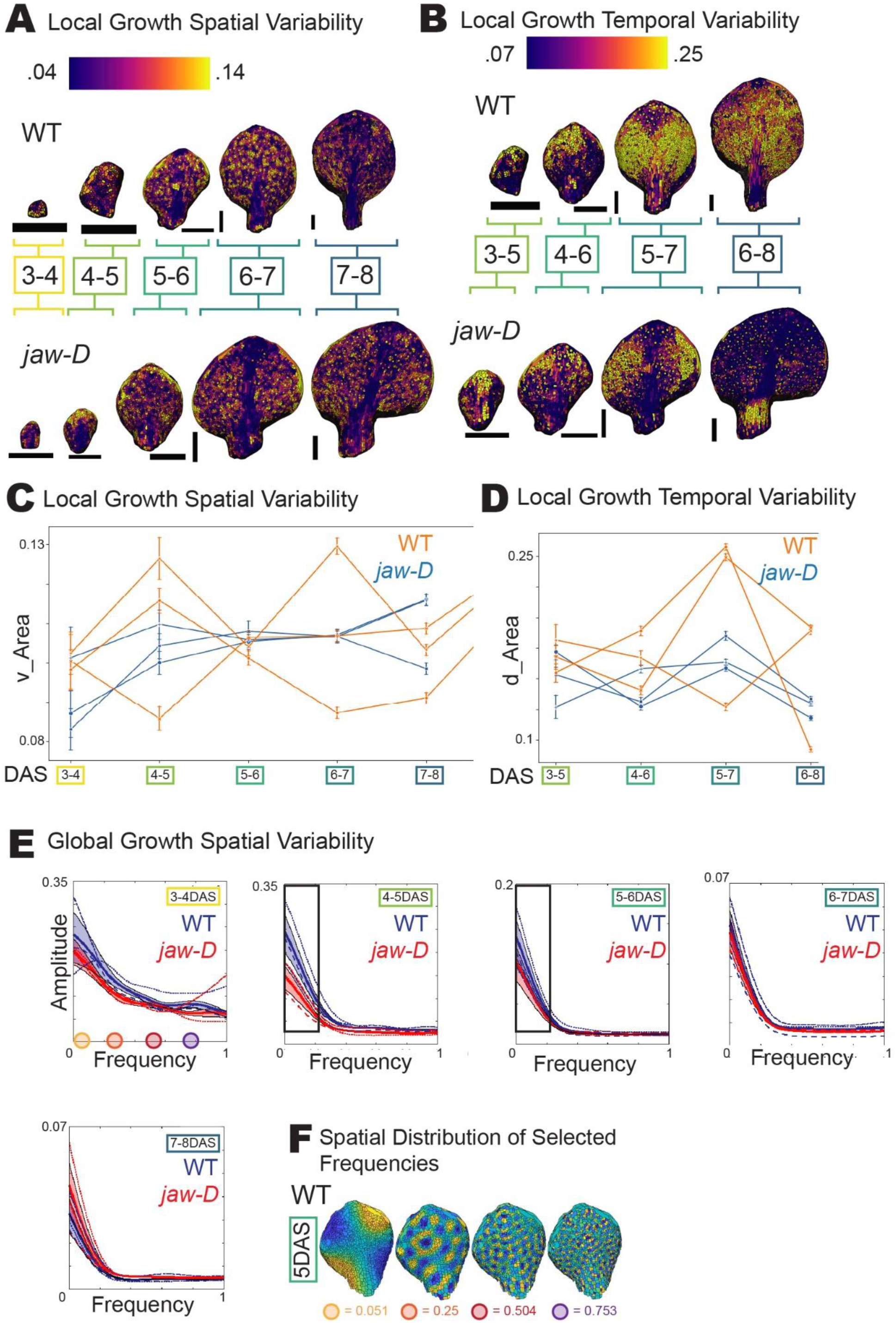
Global growth variability dominates in WT leaves. (A) Heat-maps of local spatial growth variability over 24 hour windows represented on sample WT (top) and *jaw-D* (bottom) leaf meshes on the earlier time step. Gold indicates high spatial variability and purple low spatial variability. See materials and methods for spatial variability calculation. (B) Heat-maps of local temporal growth variability in 48 hour windows, represented on sample WT (top) and *jaw-D* (bottom) leaf meshes on the middle time step. Gold indicates high temporal variability and purple low temporal variability. See methods for calculation of temporal variability. Black scale bars = 200 um. (C-D) Quantification of the mean (C) local growth spatial variability or (D) local growth temporal variability for time windows as in (A-B). Individual lines connect mean values for each of three WT (orange) or *jaw-D* (blue) replicates. Error bars indicate standard error of the mean. WT and *jaw-D* local spatial and temporal growth variability are largely indistinguishable, though *jaw-D* replicates are generally tighter around the mean. (E) Global spatial variability of areal growth for 24 hour windows determined through cellular Fourier analysis. High amplitudes at lower frequencies represent higher variability at larger length scales. Lines represent each of three WT (blue) or *jaw-D* (red) replicates with root mean squared error shaded. At 4-5DAS and 5-6DAS WT leaves exhibit much higher variability in growth across the organ than *jaw-D* (boxed). (F) Example of spatial range for different frequencies displayed on 5DAS WT leaf. Colored dots indicate the frequency value on the x-axis of (E) panel 1.

To examine larger patterns of growth change across the organ, we next applied a cellular Fourier analysis, which quantifies the heterogeneity in growth of the tissue across all length scales [52,53]. We found that there was less large-scale variation in growth rate across the organ in *jaw-D* than wild type (Fig 5E-F). This analysis verified our observations that *jaw-D* exhibits a flattened basipetal growth gradient and thus more homogeneous growth in the proximal distal axis of the leaf. Our cellular Fourier analysis also emphasized that the globally heterogeneous growth present in wild type and reduced in *jaw-D* is prevalent at 4-6, when the curvature in *jaw-D* is established (Fig 5E black boxes). Overall, our growth variability analysis confirms the two major growth defects in *jaw-D* leaves relative to wild type.

### Clonal analysis reveals contribution of progenitor cells to the leaf

To understand the extent to which each progenitor cell at 4 DAS contributes to the final leaf at 8 DAS, we analyzed the clonal sectors: the set of daughter cells at 8 DAS derived from a single progenitor cell at 4 DAS based on our lineage tracking data (Videos 1-4). Given the basipetal growth gradient in wild type, we expected cells near the distal tip of the 4 DAS leaf to differentiate early and contribute relatively little to the leaf blade, whereas cells at the proximal base would be expected to differentiate later and contribute a much larger fraction of the leaf.

Progenitor cells at 4DAS were assigned into 10 even proximal-distal bins based on their position (Fig 6A-B). Then their contributions to the final leaf at 8 DAS were compared. As expected, cells from the wild-type proximal regions at 4 DAS contributed more to 8 DAS leaves in wild type (large region of the leaf filled with dark purple in Fig 6A) than cells from the distal tip (very small region of light purple in Fig 6A), confirming the basipetal growth gradient. In contrast, in *jaw-D*, there was a more even contribution of progenitor cells along the proximal distal axis to the final leaf (more equal stripes of each purple hue in Fig 6A). We further quantified this difference by comparing the relative location of the cell along the proximal distal axis of the leaf at 4 DAS (i.e. 1.0 is the distal tip and 0.9 is 90% of the distance along the leaf) with the relative location of the clone at 8 DAS (Fig 6B). The most proximal cells at 4 DAS contribute to about 30% of the wild-type leaf whereas only about 15% of the *jaw-D* leaf (Fig 6B). Our results are consistent with the flattening of the basipetal growth gradient in *jaw-D* such that most regions of the leaf give rise to more equal amounts of leaf tissue. On the other hand, the most distal two tenths of both wild-type and *jaw-D* leaves had the same mean position from 4 DAS to 8 DAS, suggesting that tip of the leaf differentiates close to 4 DAS and may be unperturbed in *jaw-D.* We similarly examined the contributions of cells along the medial-lateral axis of the leaf and found that j*aw-D* showed a slight shift in the contribution of more lateral cells to the final leaf width (Fig 6C-D), which is consistent with the slow growing midrib region. Note, we also performed this analysis from 3 DAS to 8 DAS, but so much of the final leaf is contributed from the base in wild type that only about half of the leaf could be back-tracked (Supplementary Fig 12). The proximal-distal patterns still largely held, however the medial-lateral differences were much less pronounced.

**Fig 6.**
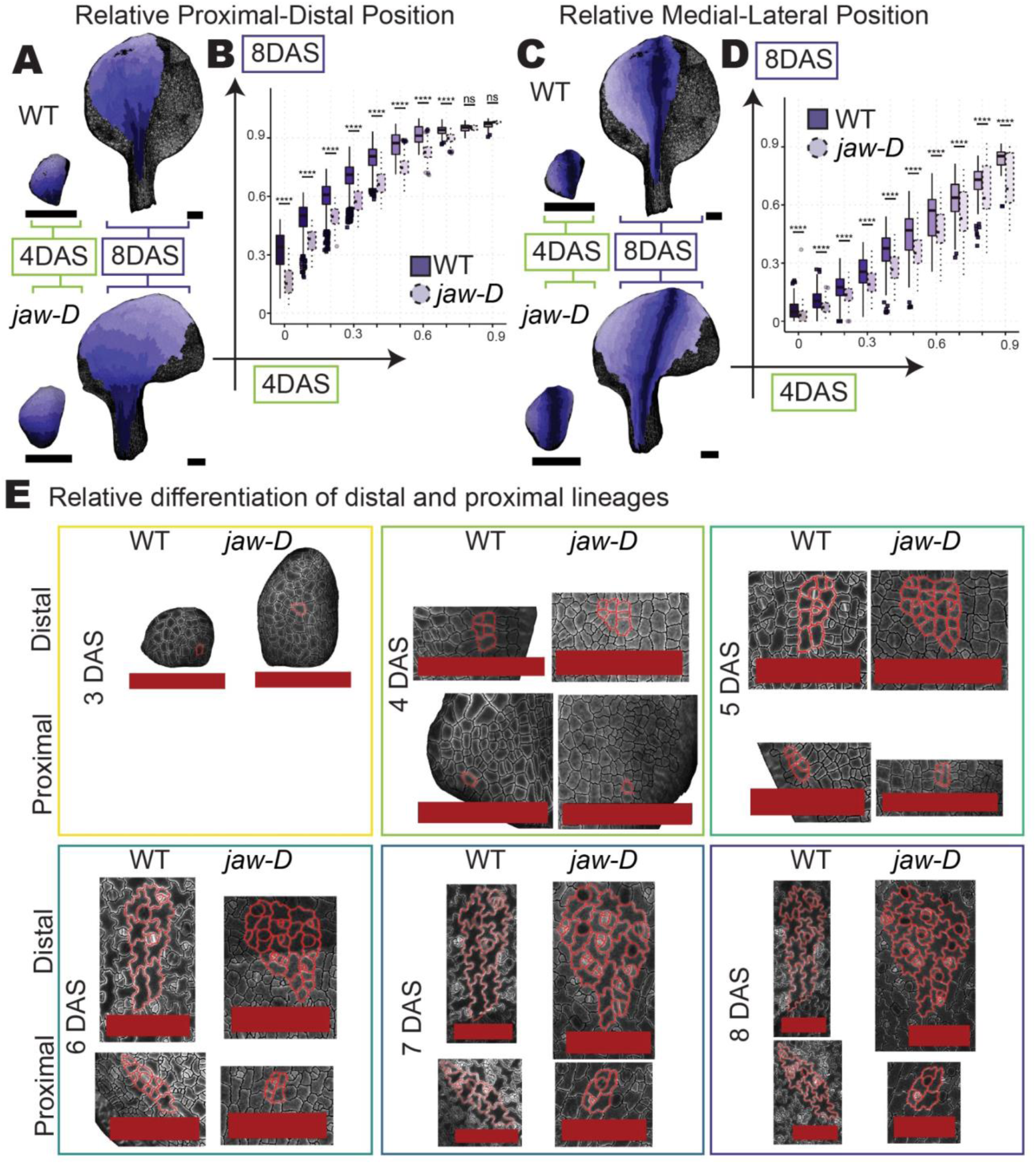
Cell differentiation quickly propagates from the distal tip along the leaf length in WT, unlike *jaw-D*. (A, C) Clonal analysis of WT (top) and *jaw-D* (bottom) cells mapped from 4 DAS to 8 DAS on sample leaf meshes. Clonal sectors (shades of purple) were binned evenly at 4 DAS into ten proximal-distal (A) or medial-lateral (C) sectors. Uncolored cells could not be tracked between all time points. (A) Darker purple WT bins end up larger than lighter bins indicative of expanded contribution of proximal growth and divisions to the overall leaf versus a more even distribution of colors in *jaw-D.* (C) Lighter purple bins are expanded in *jaw-D* indicating a lateral shift of sector contributions, . Black scale bars = 200 µm. (B,D) Quantification of proximal-distal (B) or medial-lateral (D) sector relationships for 4 DAS sector bins mapped to 8 DAS positions for all three replicates of WT (left, dark shading, solid lines) and *jaw-D* (right, light shading, dotted lines) whole leaves. Boxes indicate the 75% interquartile range, middle bar represents median, whiskers represent outliers and squares (WT) or circles (*jaw-D)* represent extreme values. Nearly all 4 DAS sector bins have different mapping to 8 DAS positions between WT and *jaw-D* representing more prominent contributions of WT proximal and medial regions than disbursed and lateral contributions in *jaw-D* (p<.0001, paired Student’s t-test and Bonferroni correction). (E) Tracing the same proximal (bottom) and distal (top) cell lineages (red outlines) for the entire experiment for WT (left) or *jaw-D* (right) sample leaves. WT cells exhibit greater differentiation into lobed pavement cells starting at 5 DAS. Red scale bars = 100 µm.

### Proximal-distal tissue differentiation corresponds with the proximal-distal gradient of growth in wild type, differentiation is correspondingly slowed in *jaw-D*

To understand how these sectoral patterns relate to cell differentiation, we used cell lobing as a measure of the differentiation of pavement cells [54,55]. We chose two lineages with approximately equal beginning and ending positions between wild type and *jaw-D* to track over the entire experiment, a distal lineage that emerged at 3 DAS and a more proximal lineage that emerged at 4 DAS (Fig 6E). In wild-type leaves, pavement cell lobing increased starting at 5 DAS in the distal lineage and 6 DAS in the proximal lineage. Lobing in the *jaw-D* lineage began at 6 DAS, never became as elaborate as wild-type lobing, and further, was more similar between distal and proximal lineages. Our results in these lineages suggest delayed differentiation of pavement cells in *jaw-D* mutants, which is consistent with the delayed progression of the basipetal wave of maturation.

The differences in pavement cell lobing observed in single lineages motivated us to measure global changes in the distribution of cellular differentiation relative to the growth front we measured in wild type versus *jaw-D.* We measured pavement cell lobing and stomatal characteristics to understand how two different types of differentiation proceed in the cells of wild-type and *jaw-D* leaves (Fig 7, Supplemental Fig S13-S14). The lobeyness measure in MorphoGraphX allows us to ascertain the level of lobing present in segmented cells throughout the epidermis of a given leaf. We used this measure to ask how patterns of differentiation present along the proximal-distal and medial-lateral axes change with leaf development. We found that in wild-type leaves, lobing begins to increase in the distal quarter of the leaf at about 5 DAS (Fig 7A; Supplemental Fig S13A). Over subsequent days, the lobeyness of many of these distal cells continues to increase and lobeyness initiation propagates from tip to base. (Fig 7A; Supplemental Fig S13A). In contrast, *jaw-D* leaf cells initiate lobing 6 DAS or later, to a lesser extent and more evenly along the length of the leaf, suggesting differentiation is delayed and reduced in *jaw-D*, consistent with the flattened basipetal growth gradient (Fig 7B, Supplemental Fig S13B). Along the medial-lateral axis, the differences in absolute magnitude of cell lobing are especially clear between wild-type and *jaw-D* samples (Supplemental Fig S13C-D). This suggests that important differentiation signals at 5 DAS that are perceived in wild type, perhaps to reinforce and structure the growth directions, are lacking in *jaw-D*.

**Fig 7.**
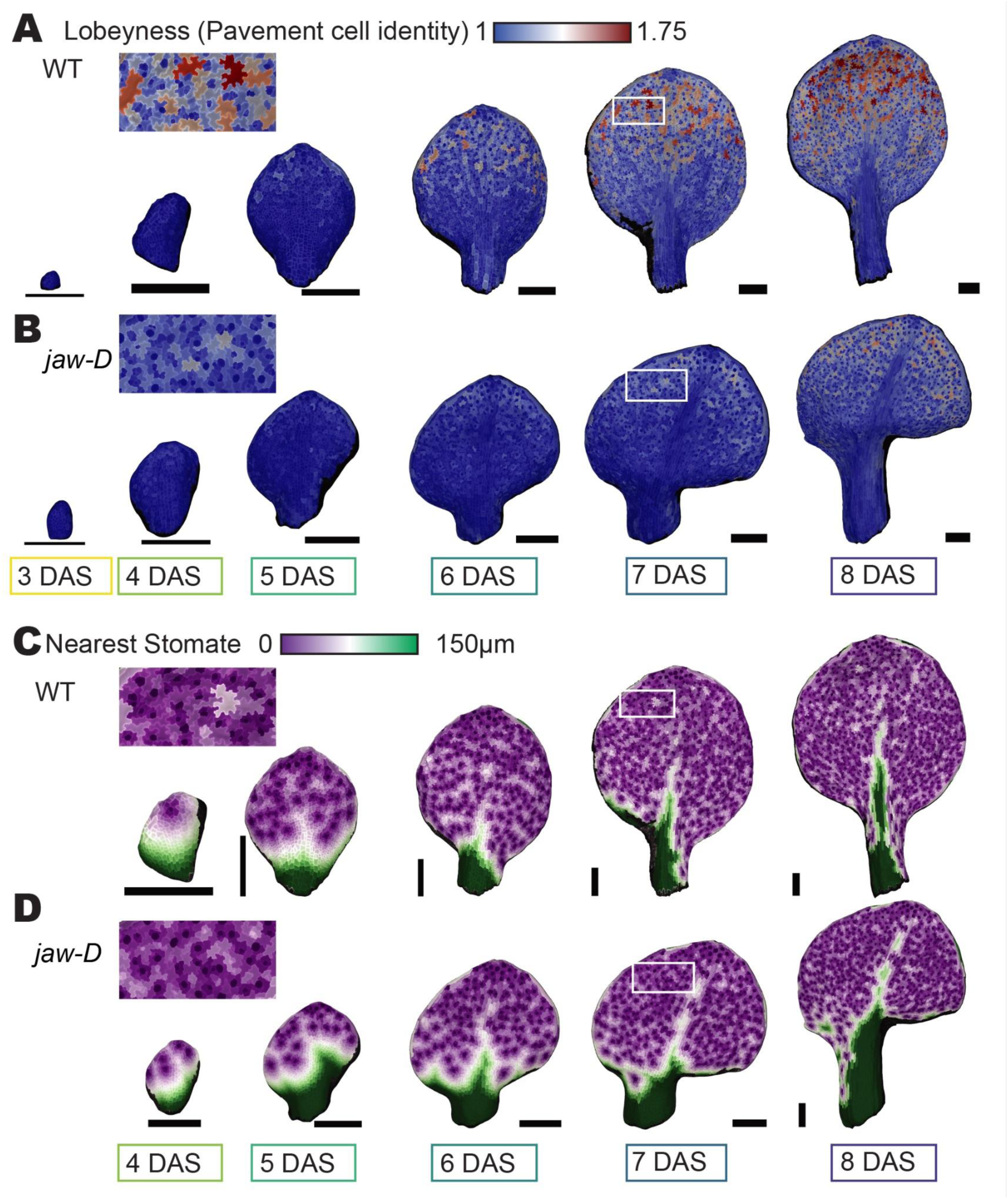
Proximal-distal tissue differentiation corresponds with the proximal-distal gradient of growth in WT, both disrupted in *jaw-D*. (A-B) Cell lobeyness (the perimeter of a cell divided by its convex hull, a measure of pavement cell differentiation). Represented as heatmaps on sample WT (A) or *jaw-D* (B) leaf meshes for each day of the imaging experiment. Black scale bars = 200 µm. Insets show magnified lobing from 7 DAS leaf (white boxes). (C-D) Stomatal spacing (the distance in µm to the nearest stomate from any pavement cell, the inverse of stomatal density) represented as heatmaps on sample WT (C) or *jaw-D* (D) meshes for each day of the imaging experiment. Insets show magnified stomatal differentiation for 7 DAS leaf (white boxes).

Stomata also initiate as leaf tissue matures, so we investigated whether stomatal production was also impaired in *jaw-D.* In contrast to pavement cell lobing, we did not see such stark differences between wild type and *jaw-D* in stomatal density (Fig 7C-D). We measured stomatal density by labeling each stomate (labeled as one cell) in each leaf and then calculated the distance from any pavement cell to the nearest stomate. At 4 DAS the spacing curve is nearly the same between wild type and *jaw-D* as the first stomata are initiated at the tip (Supplemental Fig S14). There is a slight lag in the flattening of this curve in *jaw-D* as stomatal initiation waves down the growing lamina at 5 and 6 DAS. Yet, by 7 and 8 DAS both wild type and *jaw-D* have evenly spaced stomata (Supplemental Fig S14). We hypothesize this indicates the independence of the stomatal patterning process from tissue and organ-wide shape regulation.

Thus, the delay of pavement cell differentiation in *jaw-D* is one more manifestation of the flattening of the basipetal wave of maturation in the mutant, and its failure to progress down the leaf. In contrast, stomatal patterning appears to be somewhat independent of this wave given that it progresses fairly normally in the *jaw-D* mutant.

### Longitudinal growth is coordinated across the width of the leaf blade to prevent growth conflicts, allowing flat development of the blade

Cells within the leaf that grow at different rates create growth conflicts, which lead to out of plane bending and curvature [18,22]. To assess the directional growth conflicts in wild-type and *jaw-D* growing leaves, we examined the cell growth rates along the proximal-distal direction and medial-lateral direction separately. We first examined proximal-distal growth at day 3-4 when the *jaw-D* leaf starts to curve. Given the basipetal growth gradient in wild type, we expected cells at the tip of the leaf to have a slower growth rate in the proximal distal axis than cells near the base of the leaf, which was confirmed in our measurements (Fig 8A). In wild type, we further see that at each height (point along the proximal distal axis), the growth rates are fairly uniform across the width (medial-lateral axis) of the leaf (Fig 8A). We reasoned that the uniformity across the width would mean that this section of leaf would extend at the same rate, and therefore, would not create a growth conflict, allowing the leaf to remain flat. Examining the leaf from the edge confirmed that the wild-type leaf is flat (Fig 8B). Performing the same analysis of proximal-distal growth rates in *jaw-D*, revealed regions of faster and slower growth across the width of the leaf at each height (Fig 8C). Particularly near the center of the leaf, the proximal-distal growth rates tended to be lower (Fig 8C), in keeping with the slow cell area growth rates observed in the midrib (Fig 3B). The *jaw-D* leaf starts to curve at the same time these uneven growth rates occur (Fig 8D).

**Fig. 8.**
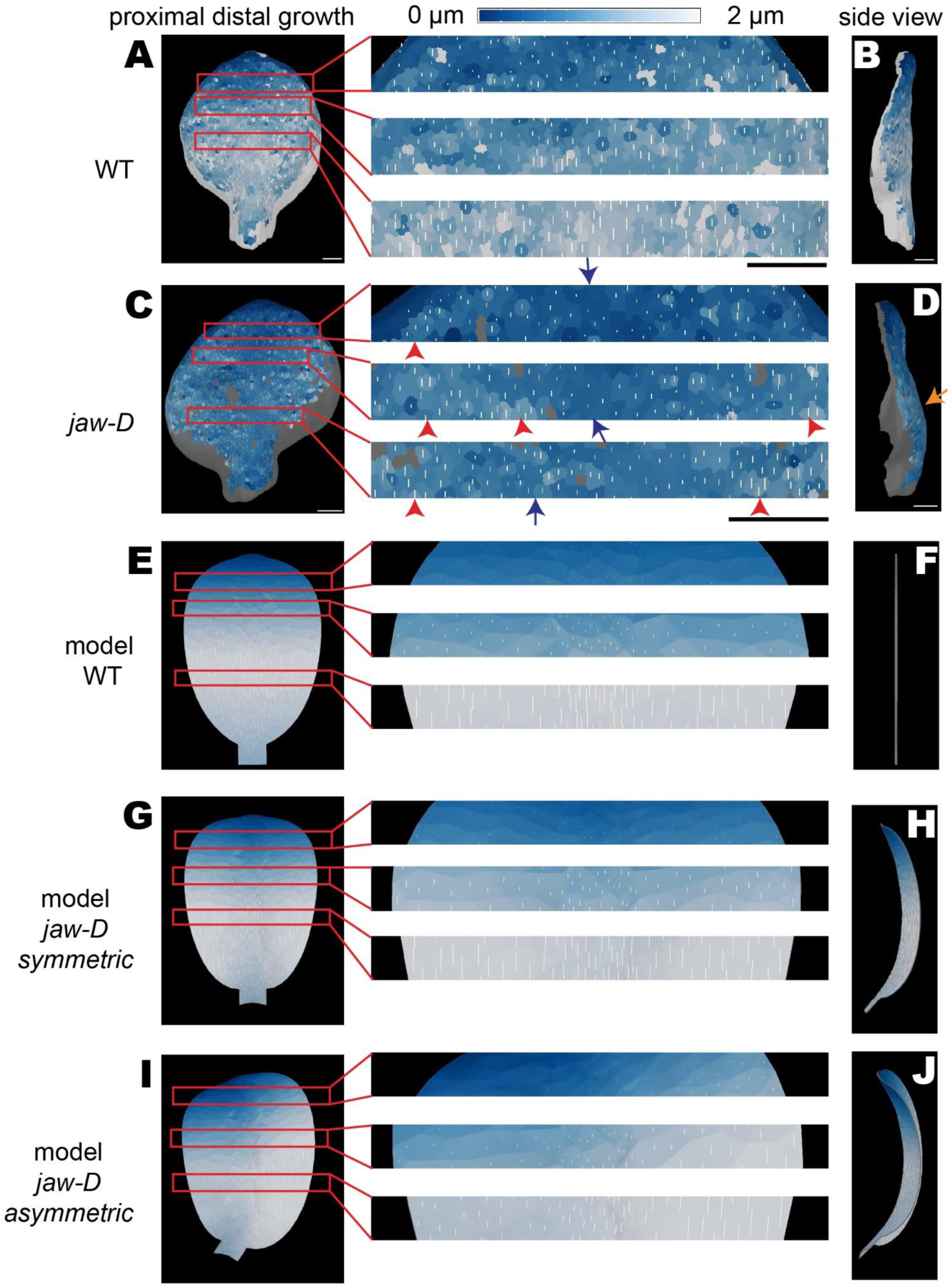
Coordination of growth rate across the width of the leaf blade prevents growth conflicts in wild type, keeping the leaf flat. (A-D) Heat map of cell growth along the proximal-distal axis of the leaf over 24 hours from 3-4 DAS in WT (A-B) and *jaw-D* (C-D). High proximal-distal extension is colored white (2 µm) and no extension is colored blue (0 µm). The magnitude of growth along this proximal-distal axis is also represented by the length of the white line drawn in each cell. This stage of leaf growth was chosen for analysis because it immediately precedes the generation of high amounts of curvature in the *jaw-D* leaf. Three regions are magnified to make the cellular growth rates visible across the medial lateral axis of the leaf. Note that the proximal distal growth rates at each medial-lateral region of the leaf are similar (allowing for typical growth variability) all the way across the leaf in WT, but not in *jaw-D* where coordination of the the proximal distal growth is disrupted and there are patches of faster (red arrows) and slower (blue arrows) growing cells. Note also the asymmetry between different sides of the *jaw-D* leaf. (B, D) A 90 degree rotation of the same leaf (side view), showing the flatness or curvature (orange arrow) of the leaf blade. (E-J) Finite element model simulation of a leaf growth in which the proximal distal growth is displayed. (E-F) When proximal-distal growth rates are equal across the width of the leaf, the leaf grows to a flat shape, similar to wild type leaves. (G-H) When the growth of the midrib is less than the blades, a growth conflict causes the leaf to curve similar to *jaw-D* leaves. (I-J) When the growth of the midrib is lower than the blades and the growth of the blades are asymmetric, the leaf also curves in asymmetric shapes similar to some *jaw-D* leaves.

We used finite element modeling to test how uniform and nonuniform patterns of proximal-distal growth affect the curvature of the leaf. We first created a model with a basipetal growth gradient in which the proximal-distal growth rate was uniform across the width of the leaf, representing wild type (Fig 8E, Video S6). In the simulation of this model, each region of the leaf grew different amounts, but since the amount was equal across the width of the leaf, it did not create any growth conflict and the leaf remained perfectly flat, similar to wild type (Fig 8F). In contrast, when we altered the proximal distal growth rate, so that it was non-uniform across the leaf, growth conflicts were created and the leaf curved similar to the large region of positive curvature at the base of the *jaw-D* leaf (Fig 8G-H, Video S7). Unlike the model, the real *jaw-D* leaf was grown between the mechanical constraints of a cover slip and the agar growth media, which is likely to cause the flattening of the tip of the leaf. We observed that *jaw-D* leaves that were not constrained in this way tended to roll back on themselves, preventing imaging, but consistent with the model. We examined *jaw-D*-like models in which the non-uniform growth across the medial-lateral axis was symmetric or asymmetric, both of which generated curvature (Fig 8I-J, Video S8). These simulations captured the variations in the *jaw-D* leaves, which range in their amount of asymmetry under our constrained growth conditions (Supplemental Fig 4-6). Thus, in wild type, within the basipetal growth gradient, growth is carefully coordinated across the medial-lateral axis to avoid growth conflict such that the leaf remains flat. Conversely, lack of coordination across the medial-lateral axis in *jaw-D* leads to curvature.

We next tested growth rates along the medial-lateral axis. Overall we see the same trend in wild type. The midrib has lower growth in width than the surrounding blade, but since the midrib has this slower rate along the whole proximal distal axis, this does not create a growth conflict in wild type. In the blade, the growth rate increases from the tip to the base due to the basipetal growth gradient (Fig S15A). This non-uniform growth rate suggests that growth conflicts are present in this direction. However, when this growth gradient in the medial-lateral axis was simulated with the model, it did not cause much effect on leaf curvature, likely because of the underlying mechanical anisotropy in the leaf tissue (Fig S15B-C, Video S9). Thus, it appears that the leaf accommodates more growth conflicts in the medial-lateral growth rates, and it is growth conflicts in proximal-distal growth that account for generation of curvature in *jaw-D*.

## Discussion

With this work, we show that coordination of growth rates across the width of wild-type leaves prevents the formation of growth conflicts, allowing the leaves to develop flat blades (Fig 9A,C). Live imaging and modeling reveal that disruption of this growth coordination across the width of the *jaw-D* leaf, particularly the slower growth of the midrib relative to the blade, causes *jaw-D* leaves to curve (Fig 9B,F). By conducting live cellular imaging on the same leaves from first initiation through a 400-fold increase in area and 100-fold increase in cell number over six days, we have captured an immense level of detail of this process (Fig 1). We found that disruptions in tissue curvature in *jaw-D* become strong at 6 DAS (Fig 2). The *jaw-D* leaves exhibit two major disruptions to the typical wild-type pattern of cellular growth. First, whereas wild type exhibits a basipetal growth gradient with a proximally localized growth maximum and slowing growth gradient towards the tip, the *jaw-D* leaf exhibits a pattern of medium growth broadly extending along most of the length of the leaf, thus flattening the basipetal gradient (Fig 3-4). Corresponding with the basipetal growth gradient, there is a basipetal wave of cell differentiation as shown by the lobing of pavement cells, which is delayed and reduced in *jaw-D* (Fig 6-7). Second, the wild-type growth rate is relatively uniform across the width of the leaf at each height, whereas the *jaw-D* leaf exhibits slower growth at the midrib than the leaf blade (Fig 3-4, 8). Thus, the wild type avoids growth conflicts, whereas they are created in *jaw-D*. Finally, we used modeling to test whether the slower growth of the midrib from the blade in *jaw-D* leaves generates the curvature of the leaf (Fig 8-9). Simulations of this growth change reveal that uniform growth rates across the width of the leaf are essential for maintaining flatness in wild type and unequal growth rates between midrib and blade generates a positive curvature in *jaw-D* (Fig 8-9). Overall, we conclude that cell growth is coordinated across the width of the wild-type leaf to maintain flatness of the leaf blade.

**Fig 9.**
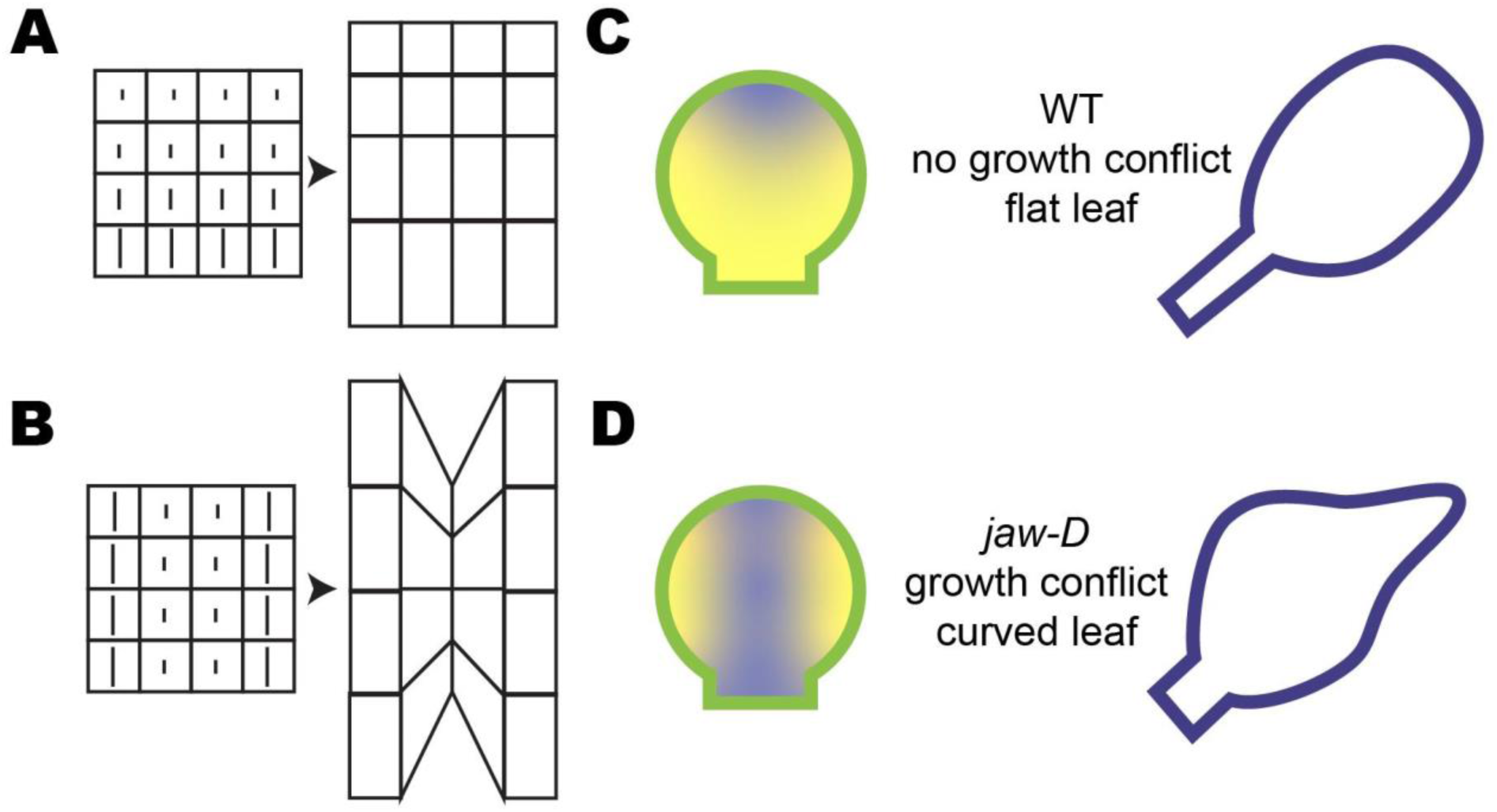
Cell growth is coordinated across the medial lateral axis of the leaf to maintain leaf flatness in WT while medial lateral growth conflicts lead to curvature in *jaw-D*. (A) Conceptual diagram illustrating that as long as growth rates are coordinated all the way across the width of the tissue, no growth conflicts are created, despite a basipetal growth gradient with slower growing cells at the top. (B) Conceptual diagram illustrating that different cell growth rates across the width of the leaf create growth conflicts that will lead to curvature. (C) The basipetal growth gradient where slower growth (purple) starts at the tip of the leaf and descends maintains coordination of growth rates across the width of the leaf and thus produces flat leaves. (D) The slower growth of the midrib (purple) than the leaf blade (yellow) in *jaw-D* creates growth conflicts causing the leaf to curve. In addition, the basipetal growth gradient is flattened in *jaw-D*.

The alterations in growth of the *jaw-D* mutant are more complex and nuanced than expected based on current understanding of the molecular function of *miR319*. The *jaw-D* mutant phenotype is caused by the insertion of strong 35S enhancer elements upstream of miR319, causing stronger expression of miR319[28] . In wild type, miR319 is expressed from the base of leaves and floral organs [41] where it targets the mRNAs encoding class II TCP transcription factors, limiting their accumulation to the distal tip of the leaf [28,29]. Class II TCP transcription factors promote cell differentiation and inhibit cell division, contributing to the formation of the basipetal gradient of growth and maturation in wild type. Our growth data agrees well with this model in wild type as a region of fast growth and division is present at the base of wild-type leaves (Fig 3-4). Since *jaw-D* over-expresses miR319 and should therefore further decrease the presence of TCPs in the leaf, we expected there to be an increase in the size of the fast growth and division zone seen in wild type. However, the change was more nuanced. Instead, the growth rates were more moderate in *jaw-D* and distributed further along the proximal-distal axis. *jaw-D* also affected the midrib versus blade growth rates, leading to a lack of coordination. We noted that cell divisions followed the altered growth pattern in *jaw-D,* and that their contribution to the organ ends earlier than growth. We also noticed an interesting difference between cellular differentiation programs. Namely, the basipetal differentiation of pavement cells is altered in *jaw-D,* while the initiation and spacing of stomata remains intact (Fig 7). Our results highlight the diversity of differentiation programs present in leaves and how high resolution spatial and temporal imaging can help to resolve the different contributions of these independent processes to leaf form and function.

In many organisms, mechanical instabilities can induce organ waving or folded forms during normal development [56]. Studies in kelp suggest that the water flow stresses experienced by the same species in different environments determines blade ruffling or flattening [21]. In animals, intestinal and cranial folding out of the plane have been attributed to differential growth and buckling between layers [57–61]. Work in *Drosophila* also concurs that the spatial distribution of differential growth is important in creating multiple folds of the wing disc epithelium [23]. Our model complements recent observations that spatially specific reinforcement of tissue stresses occurs in three dimensions to promote the initial flattening of the rounded leaf primordium to form a lamina [62]. Likewise, cell wall integrity sensing and proper response to mechanical signals have recently been found to be required for maintaining the flatness of the thallus in the liverwort *Marchantia polymorpha* [63].

Additionally, while many leaves from different species exhibit a similar basipetal growth gradient as Arabidopsis, leaves from some species exhibit acropetal (growth slowing progressively from base to tip), diffuse (growth throughout) or bidirectional (growth from the center of the leaf) growth patterns. The microRNA miR396, which represses growth by inhibiting GRFs, has been shown to be present in basipetal, acropetal and uniform patterns along the proximal-distal axes of different plant species [13]. It would be interesting if the growth patterns we measured had corollaries in these species where the miR396 pattern differs from Arabidopsis. We predict these cases would follow our general mechanism - that the growth across the width of the leaf at each height remains uniform, though the specific distributions of growth rate zones may vary, to ensure development of a flat blade. As our study highlights, future work on leaf flatness should focus on the long distance molecular and mechanical mechanisms coordinating cellular growth across the width of the leaf to prevent growth conflicts and maintain flatness.

## Materials and Methods

### Plant Growth Conditions

For live imaging, seeds were surface sterilized by washing once with 70% EtOH, 0.05% SDS solution for 7-10 minutes on a nutating shaker, then 3 subsequent washes with 100% EtOH, then dried on autoclaved filter paper. Seeds were sown on 0.5x Murashige and Skoog media (pH 5.7, 0.5 g/L MES, 1% phytoagar) supplemented with 1% sucrose. For genetics, plants were grown to maturity on Lambert Mix LM-111 soil. All plants were stratified for at least 2 days at 4°C then transferred to Percival growth chambers and grown under 24-hour fluorescent light, ∼100 μmol m^−2^s^−1^ at 22°C.

### Plant accessions

Wild-type plants are ecotype Col-0. *jaw-D* leaves are the *jaw-1D* allele from [28]. Plants were crossed with the epidermal specific fluorescent reporters for plasma membrane (pAR169 *AtML1:mCitrine-RCI2a*) and nucleus (pAR229 *AtML1:H2B-TFP*) [64,65]. In subsequent generations, plants homozygous for both markers were selected. These lines have been deposited in the ABRC under accession numbers CS73343 (wild type pAR169 pAR229) and CS73344 (*jaw-D* pAR169 pAR229). Note, only the plasma membrane marker was imaged for the purposes of this paper.

### Sample preparation – live imaging

Seedling germination times have inherent variability [66]. In our hands, we found that the first two leaves of wild-type and *jaw-D* plants initiated three days after being exposed to our growth conditions (reported as Days After Sowing - DAS). At this stage, the cotyledons are fully greened and have bent at 90° angles from the stem. Therefore, we dissected cotyledons off the early seedlings 2 DAS, allowed the plants to rest for one day and then proceeded to capture images of growing first leaves using lines expressing the plasma membrane marker. Coverslips were suspended a few millimeters above the sample with vacuum grease to encourage leaves to remain in good imaging positions. Samples beneath coverslips were submerged in perfluorodecalin, a high refractive index compound that allows for proper gas exchange during imaging. More detail about our imaging method can be found in our publication [47].

### Confocal imaging

Plants were imaged on a Zeiss 710 upright laser scanning confocal microscope with 20x Plan-Apochromat NA 1.0 water immersion lens, with z-step between 0.5um and 1.0um. The plasma membrane mCitrine marker was excited with a 514 nm wavelength laser at 1-2% power and the emission spectra from 519-650nm was collected. As samples grew and were too large to image in one stack, separate scans were taken and stacks manually aligned and merged in MorphoGraphX 2.0 [48].

### Fully grown leaf imaging

First and second leaves were harvested from plants once fully grown, at least 31 DAS, and scanned with a Canon CanoScan LiDE 110 scanner.

### Image processing – live imaging

Zeiss .LSM imaging files were converted to .TIF in ImageJ. Tiff files were imported into MorphoGraphX 2.0 [48]. The MorphoGraphX image processing pipeline has been described in detail [47]. Briefly, masks were created of the confocal stacks through 1-3 rounds of Gaussian blurring, then edge detection, closing holes in older samples where masks showed gaps, surface creation, manual selection and deletion of the surface that did not contain confocal signal and then projection of the confocal signal onto the surface. Meshes were subdivided once, then subject to 2-3 rounds of auto-segmentation, adaptive mesh subdivision at the new cell borders and projection of the confocal signal back onto the refined mesh. Cell segmentations were manually corrected immediately after segmentation or through the process of manual cell lineage tracing and cell junction correction using the check correspondence process. Cell and growth parameters were measured and heat maps generated using a python script run through MorphoGraphX 2.0. All >77,000 cells in the final analysis were selected based on the presence of lineage tracking ensuring that only cells that were manually curated were included in the final dataset. All segmented meshes and measures are included at https://doi.org/10.17605/OSF.IO/D7X3Y).

### Image processing - fully grown leaves

Scanned leaf images were processed using the ROI tool in ImageJ. Each leaf was first outlined with the white paintbrush tool to color over bits of dirt and shadows. Images were then converted to 8-bit grayscale and a threshold of 187 was applied. To measure leaf area, each leaf outline was selected with the wand tool, filled in using the Ctrl+F shortcut and measured with the Ctrl+M shortcut. The whole leaf and lamina length were measured using the segmented line tool by tracing a line from tip to base of the lamina and then along the middle of the petiole. The lamina was then digitally separated from the petiole by drawing over the petiole neck with the paintbrush tool. The separated lamina was selected with the wand tool and area measured with Ctrl+M. Petiole length and area were determined by subtracting the measured lamina values from the whole leaf values. Ripples were counted as protrusions visible in the thresholded images that were symmetrical along the proximal-distal axis (see Fig 1K). Bulges on one side were counted as half ripples.

### Data analysis

All data processing, analysis and plotting was performed in RStudio [67,68]. Sigmoidal models were fit using [69]. CVs were compared using [70]. Scripts used to process the data and create each figure can be found at https://github.com/kateharline/jawd-paper. We have provided the segmented meshes, measures of cellular growth, divisions and characteristics to enable further exploration into the variety of cell lineages we captured in this data set available at osf.io (https://doi.org/10.17605/OSF.IO/D7X3Y).

### One dimensional model of leaf length

We develop a one-dimensional discrete-time model to capture the leaf length dynamics for WT and *jaw-D* genotypes. Our goal was to explore how contrasting spatial growth profiles for WT and jaw-D genotypes can lead to overall similar leaf length dynamics. The model considers a spatio-temporal cellular growth rate at any position along the leaf length and over time, and uses it in a discrete-time model setting to compute the length across days. For technical details, please refer to S1 Supporting Information.

### Analysis of Local Temporal and Spatial variability in Cell Growth

To quantify local temporal and spatial variability in cell growth we follow [50]. The areal growth rate of a cell between two consecutive time points is defined by *AGR*_1_ = (*A*_1_/*A*_2_)/*t*, where *A*_1_ and *A*_2_ denote the cell area in the first and second time point, respectively, and *t* is the elapsed time between the time points measured in hours. If the cell has divided between the two time points *A*_2_ is the sum of the areas of all daughter cells. Local temporal variability in cell area growth is measured using the formula 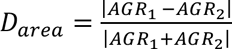. Since this measure is calculated based on 3 consecutive time points it is ambiguous for which time point we are measuring temporal variability. To resolve this ambiguity, we always use the formula to measure the temporal variability at the second (middle) time point. Given a cell *c* the local spatial variability of the cell is measured using its areal growth rate *AGR*(*c*) and the areal growth rates of its bordering neighbors *AGR*(*i*), *i* = 1,2, . . ., *k*, where *k* denotes the number of bordering neighbors. Local spatial variability in cell area growth is measured using the formula 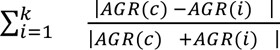. The local spatial variability is only calculated based on two time points and the formula is used to measure the variability at the first time point.

### Cellular Fourier analysis

Below we summarize the main steps of the spectral analysis. For more technical details, refer to S1 Supporting Information and [52].

We analyze the areal growth rate and the deformation of tissues. The areal growth rate of each cell is defined as the relative difference of their surface areas at successive time steps divided by the time delay between two time steps. The deformation is a symmetric matrix summarizing the two principal directions of growth, and the ratios to apply in these two directions to deform a cell’s contour at a preceding time step to the same cell’s contour at a following time step.

We then perform the spectral analysis consisting of representing the spatial variation of the areal growth or of the deformation matrix as a linear combination of harmonics. Each harmonic is associated with a frequency characterizing its spatial variation scale. By performing our analysis, we obtain the Cellular Fourier Transforms of the areal growth rate and of the deformation. These are the amplitudes with which the different harmonics decompose the areal growth rate and the deformation matrix. For a tissue, the first harmonic is a constant associated to the spatial frequency 0, and the corresponding CFT for the areal growth rate is the whole tissue’s areal growth. The two following harmonics vary at the scale of the tissue and the associated CFTs for the areal growth rate give the average growth gradient at the tissue scale. The higher the rank of the harmonics is, the higher the spatial frequency and the smaller the considered space scale are. The deformation being described by a matrix, its CFTs are also matrices. The first CFT will, for example, be the average tissue deformation. We notice that, although a cell’s deformation is 2D and lies in its tangential plane, the overall tissue deforms in 3D if it is not flat. The CFTs of the deformation matrix depend therefore on the tissue geometry. To analyze growth anisotropy, we considered the difference between the first two eigenvalues of the CFTs of deformation which we divide by the time delay between two time steps. It quantifies the deviation from isotropic deformation.

The spectra we finally considered are those of the areal growth rate and the deviation from isotropic growth. These spectra have been smoothed to be comparable among different tissues. In these spectra the y-axis represents the CFTs and the x-axis spatial frequencies. Low frequencies correspond to large spatial variation scales and high frequencies to small spatial variation scales. Spatial frequencies equal to one correspond to the cell scale (mean cell size).

### Finite element method simulations

The template for the growing leaf model was created from an initial flat 2D contour representing the young leaf shape. This was triangulated and then extruded into a sheet made of a single layer of wedges (prisms). This follows the structure of previous models of leaf shape [8], although in our model a small amount of noise was added to the vertices in the normal direction allowing the structure to bend out of the plane. The MorphoMechanX (www.MorphoMechanX.org) modeling framework was used for the FEM simulation [71–73]. An isotropic linear material model was used. Growth was implemented by increasing the size of the reference configuration of the elements, with independent control of growth parallel or perpendicular to a growth axis. The rates of growth and the growth direction were specified manually through the software interactive GUI based on qualitative assessment of live imaging data. After each growth step, the mechanical equilibrium was found, and residual stresses were released.

The mathematical details about mechanical equilibrium calculation and growth algorithm are provided together with the simulation models in S1 Supporting Information

## Acknowledgements

The authors would like to thank the members of the Roeder, Scanlon, Boudaoud, Hamant and Smith labs for helpful discussions and advice on the manuscript. We thank Javier Palatnik for information on the *jaw-D* insertion site. We thank Zahava Rojer for technical assistance.

## Author Contributions

K.H., J.W.S. and A.H.K.R conceived experiments. K.H. conducted live imaging experiments, image processing, cell growth and characteristics data analysis. K.H., A.H.K.R., A.F. A.B., N.K., C-B.L., R.S.S. conceived analyses. A.F. conducted cellular fourier analysis. N.K. conducted spatio-temporal analysis. B.L., A.S. and R.S.S. conducted leaf growth modeling. S.S. and R.S.S. developed and provided support for MorphoGraphX. G.M. and R.S.S. developed mechanical modeling software MorphoDynamX. K.H. and A.H.K.R. wrote the manuscript and assembled figures. All authors contributed to manuscript preparation and editing.

## Data availability

Live imaging datasets, models and analysis scripts are available at osf.io (https://doi.org/10.17605/OSF.IO/D7X3Y).

Scripts used to process the data and create each figure can be found at https://github.com/kateharline/jawd-paper.

Seeds expressing the fluorescent plasma membrane and nuclear marker (pAR169 mCitrine-RCI2A and pAR229 *ML1::H2B-mTFP*) in wild type (Col-0) are available from the Arabidopsis Biological Resource Center (ABRC) under stock number CS73343. Seeds expressing the fluorescent plasma membrane and nuclear marker (pAR169 mCitrine-RCI2A and pAR229 *ML1::H2B-mTFP*) in *jaw-D* are available from ABRC under stock number CS73344.

