## Supplemental Figures for "Cell growth rates coordinate across the width of the leaf to remain flat"

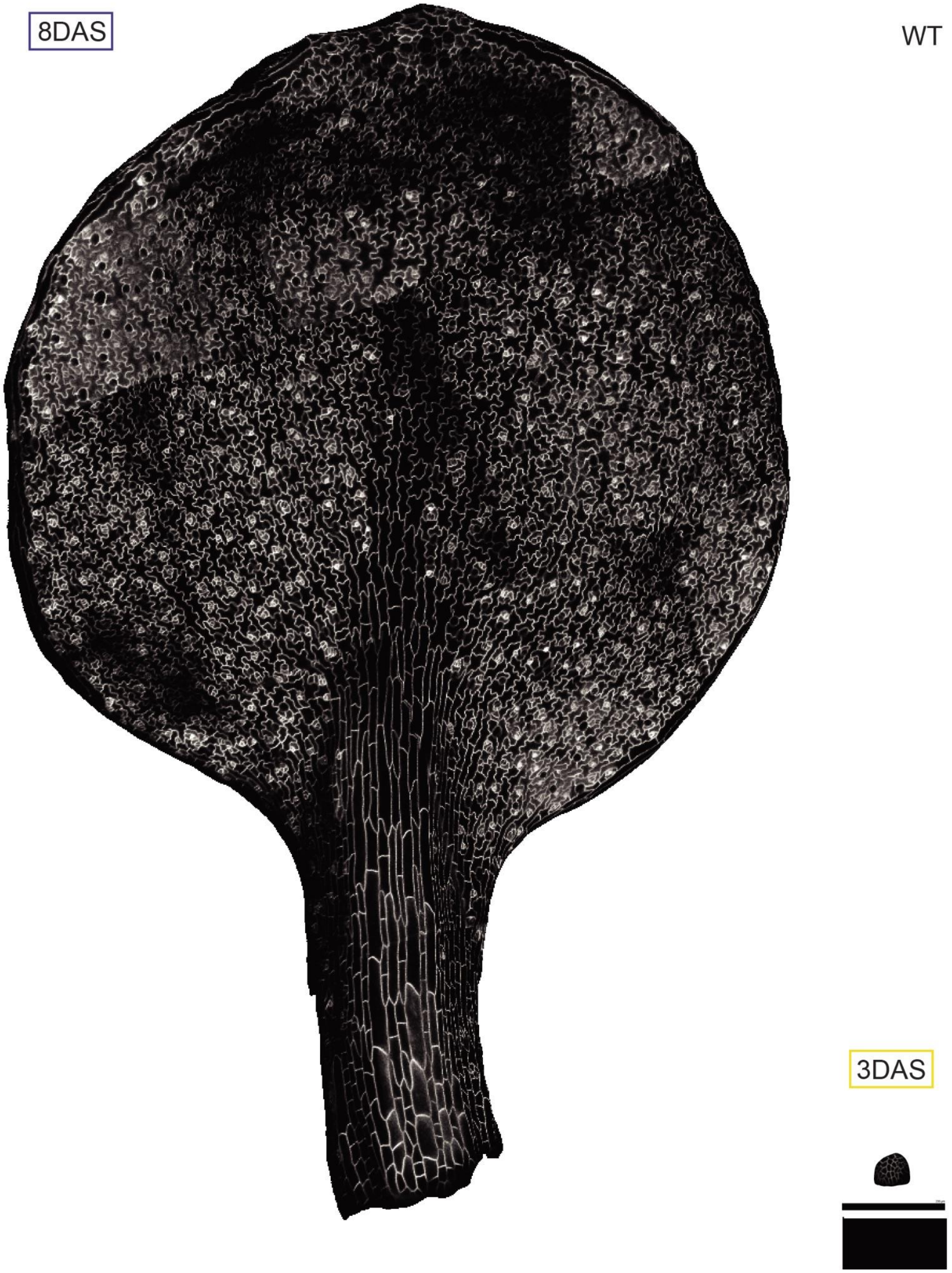


**Supplemental Fig. S1 WT leaf live imaging replicate 1**
Higher magnification image of main text 8 DAS WT leaf (left) and its corresponding, to scale, 3 DAS primordia (right). Black bar = 200 µm. Associated with Fig 1.

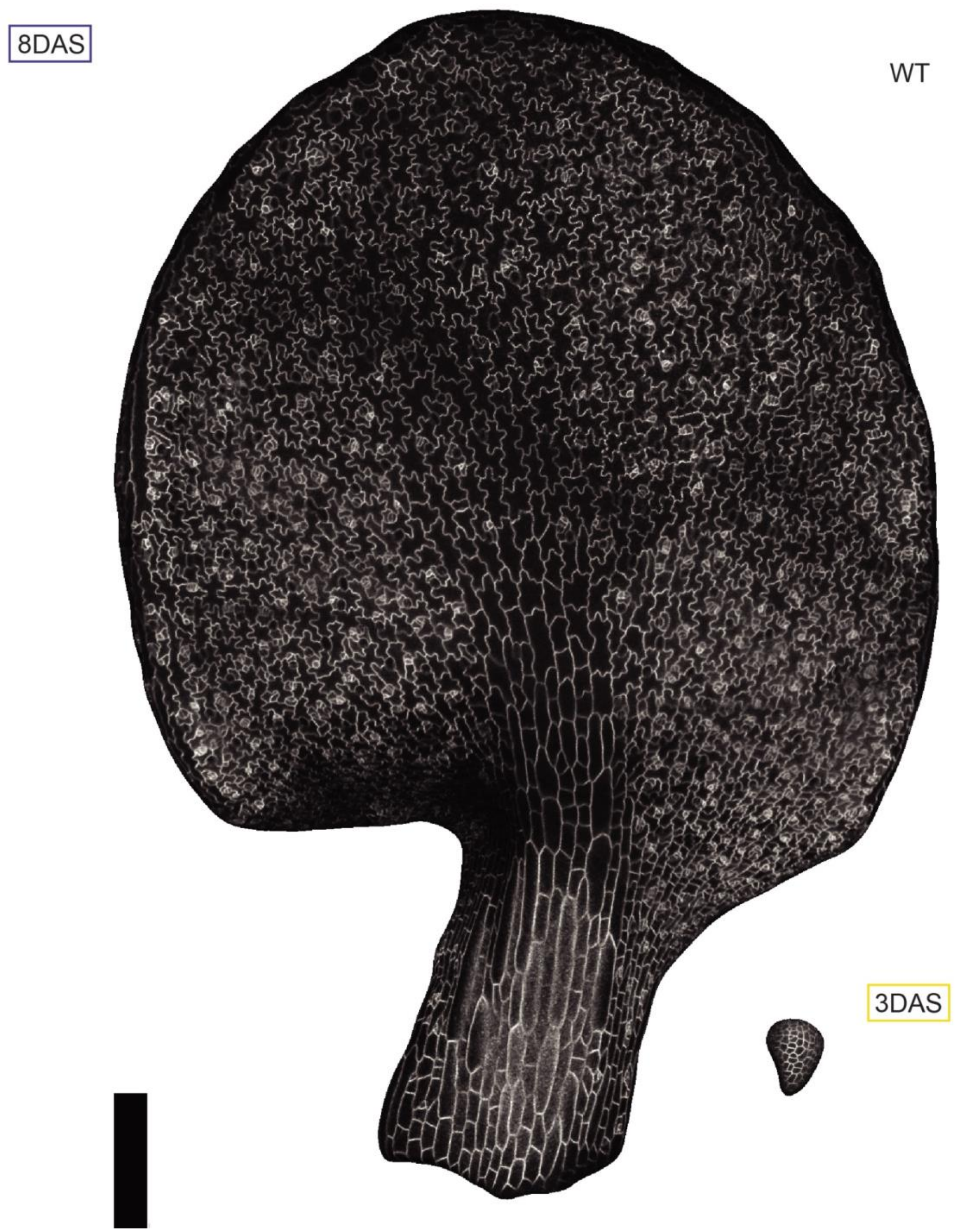


**Supplemental Fig. S2 WT leaf live imaging replicate 2**
Higher magnification image of replicate 8 DAS WT leaf (left) and its corresponding, to scale, 3 DAS primordia (right). Black bar = 200 μm. Associated with Fig 1.

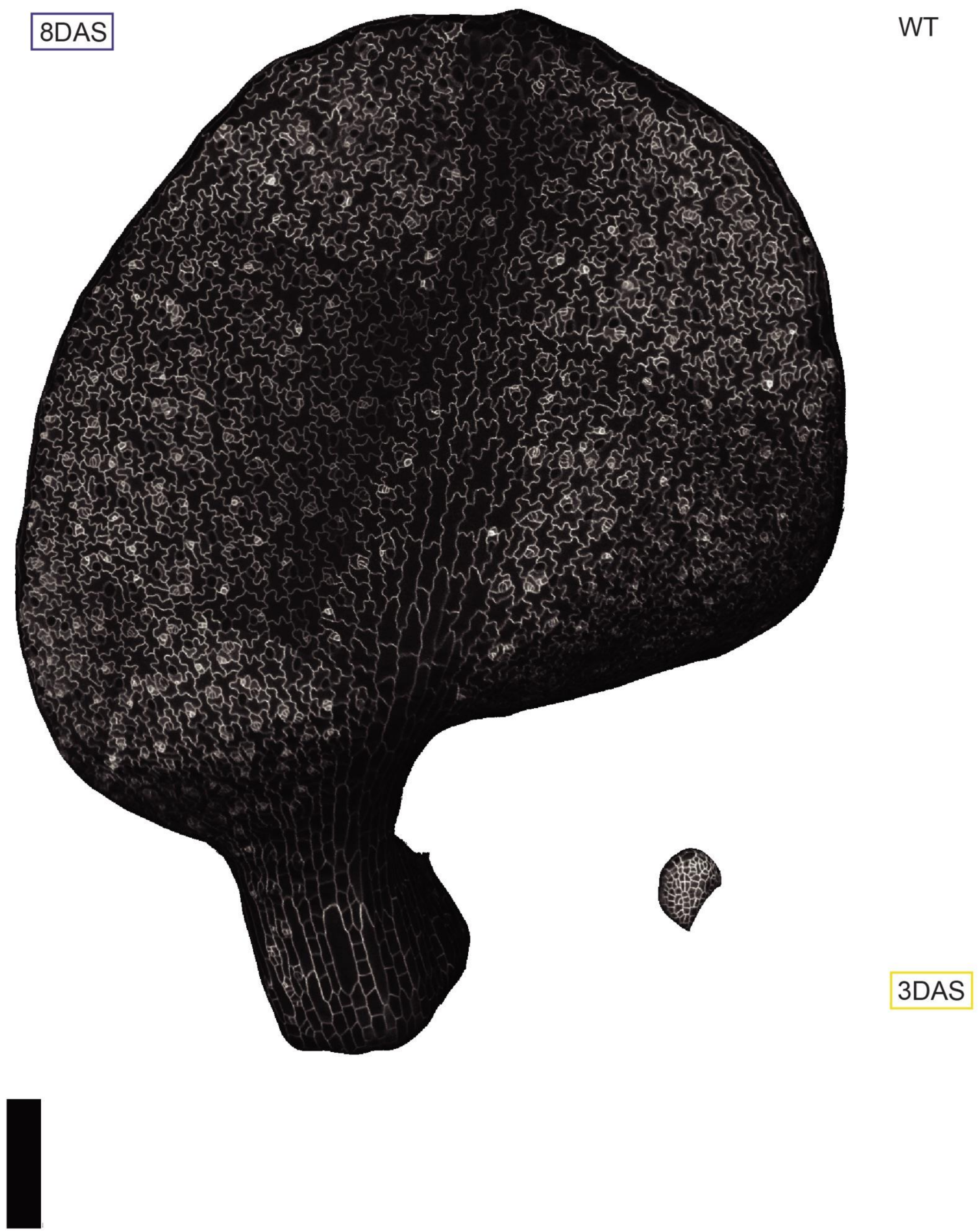


**Supplemental Fig. S3 WT leaf live imaging replicate 3**
Higher magnification image of replicate 8 DAS WT leaf (left) and its corresponding, to scale, 3 DAS primordia (right). Black bar = 200 µm. Associated with Fig 1.

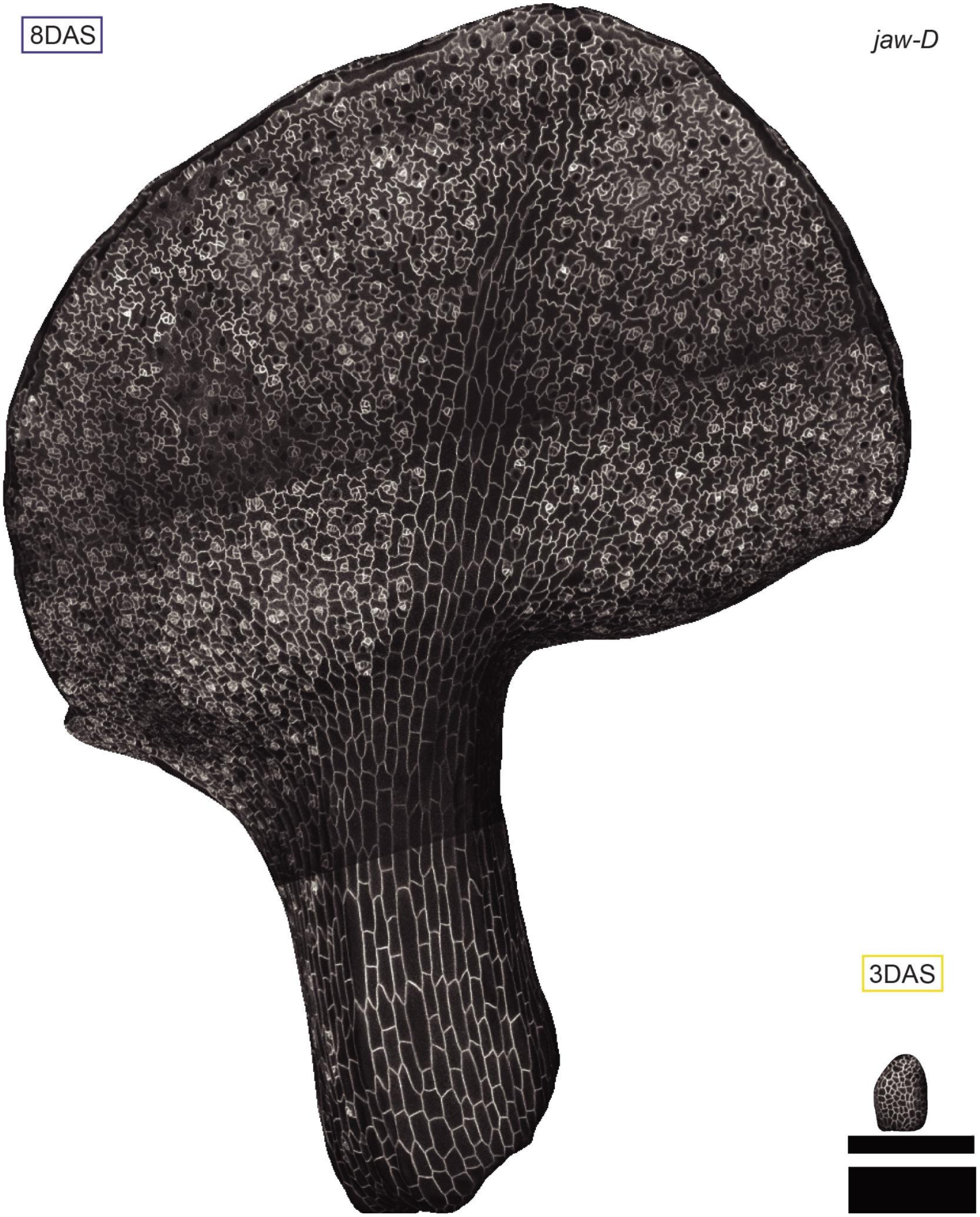


**Supplemental Fig. S4 *jaw-D* leaf live imaging replicate 1**
Higher magnification image of main text 8 DAS *jaw-D* leaf (left) and its corresponding, to scale, 3 DAS primordia (right). Black bar = 200 µm. Associated with Fig 1.

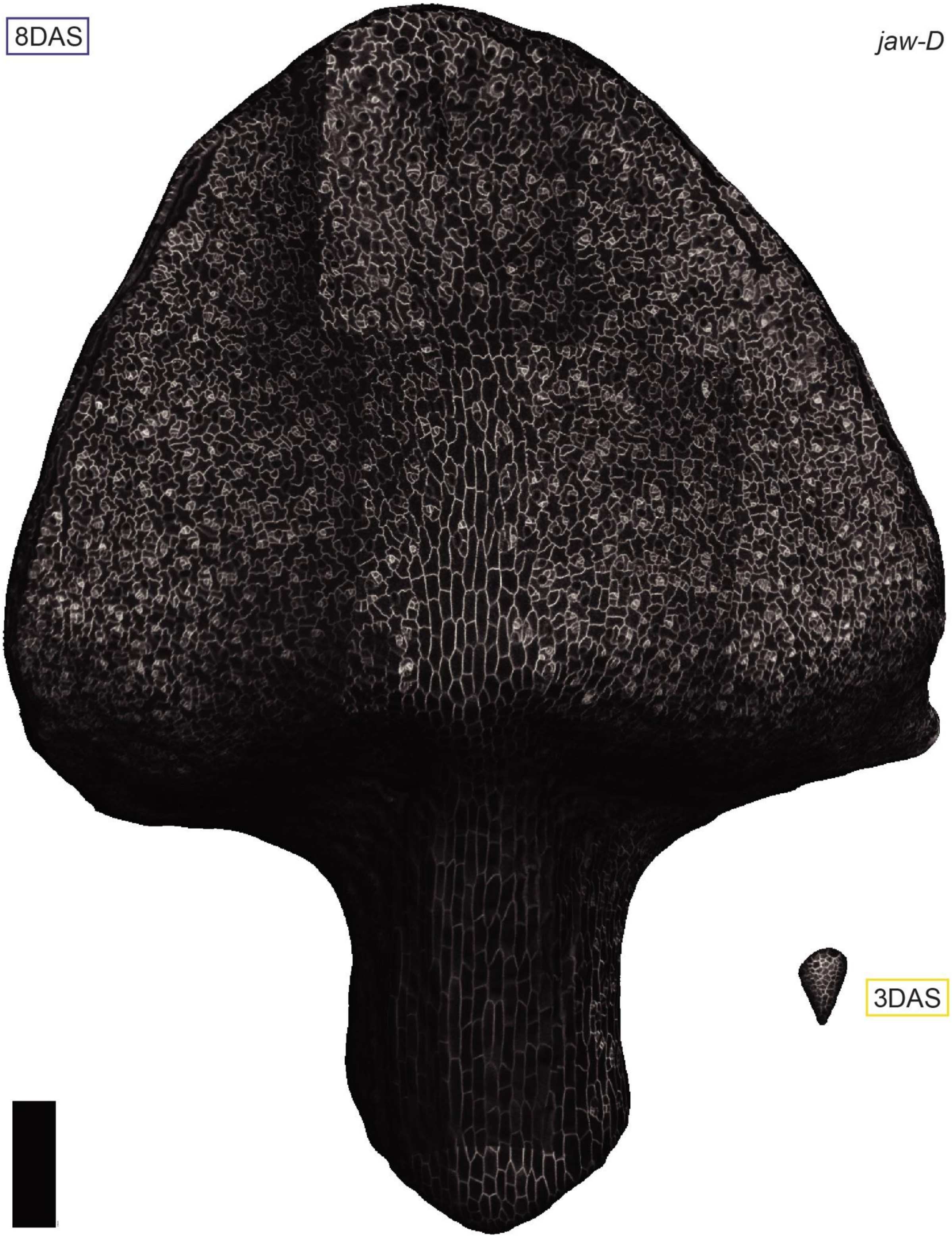


**Supplemental Fig. S5 *jaw-D* leaf live imaging replicate 2**

Higher magnification image of replicate 8 DAS *jaw-D* leaf (left) with less visual skew and its corresponding, to scale, 3 DAS primordia (right). Black bar = 200 µm. Associated with Fig 1.

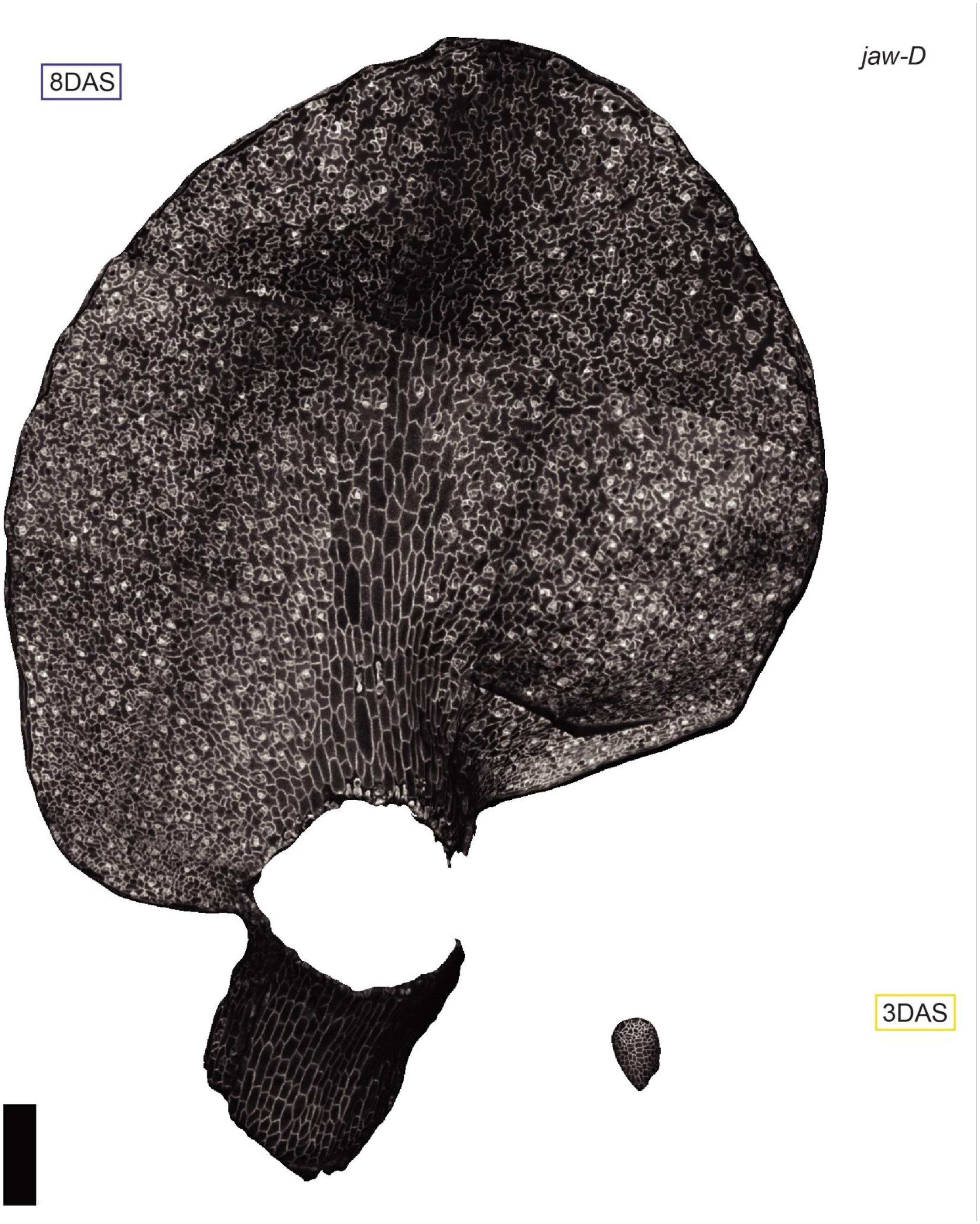


**Supplemental Fig. S6 *jaw-D* leaf live imaging replicate 3**
Higher magnification image of replicate 8 DAS *jaw-D* leaf (left) and its corresponding, to scale, 3 DAS primordia (right). Note the petiole broke due to the strong curvature. Black bar = 200 µm. Associated with Fig 1.

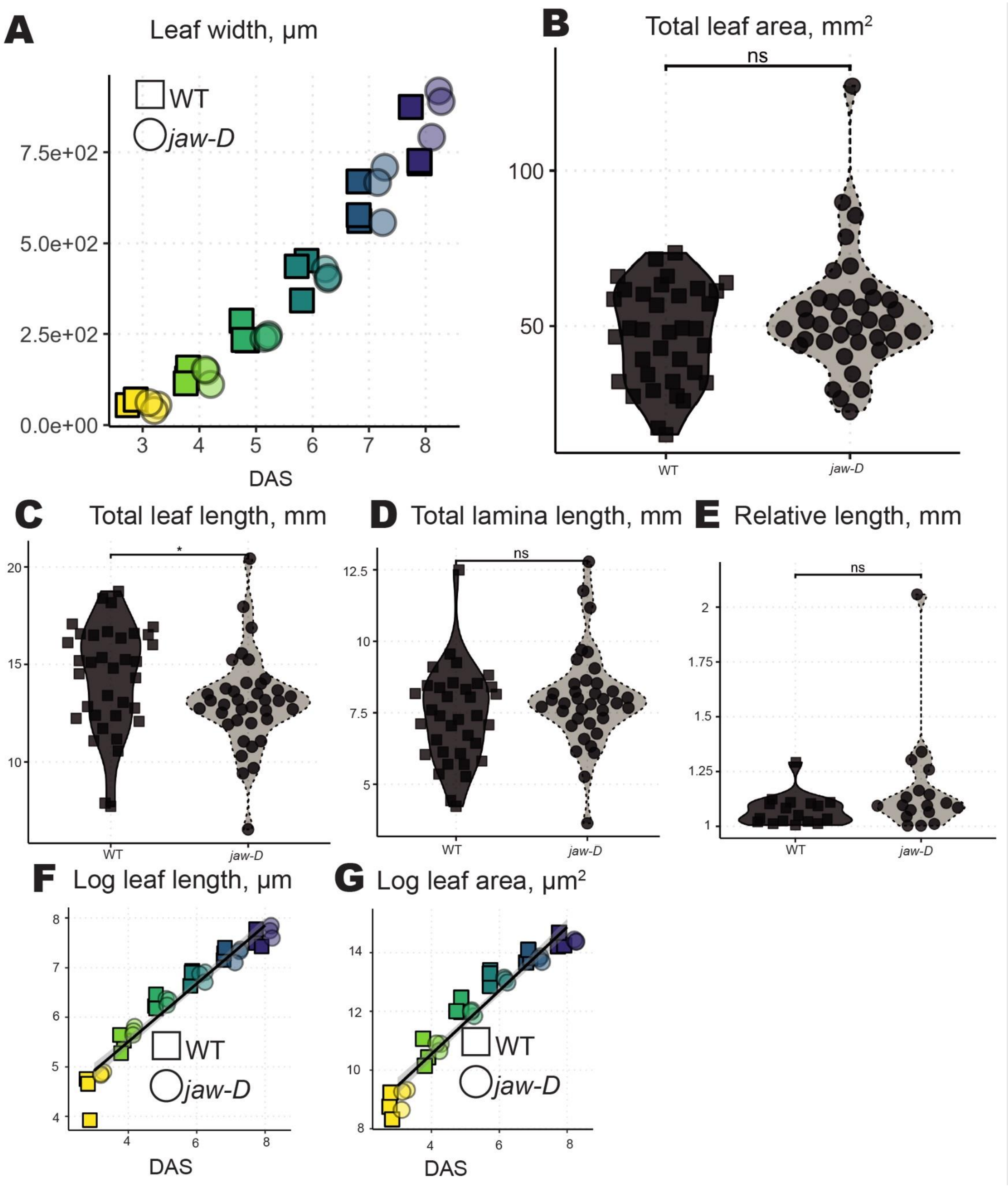


**Supplemental Fig. S7 Additional features of live imaged and fully grown leaves.**

(A) Leaf width increases exponentially in WT (squares) and *jaw-D* (circles) leaves during the live imaging experiment. Colors indicate DAS.

(B) Final leaf area of fully grown first and second leaves between WT and *jaw-D* is not significantly different, though the distributions are distinct with WT showing a bimodal distribution and *jaw-D* with a more common median value and more extreme values. Color,

shape and linetype indicate genotype (solid, squares, dark gray = WT; dotted, circles, light gray = *jaw-D*)
(C) Total leaf length of fully grown first and second leaves between WT and *jaw-D* shows a similar pattern with slightly shorter leaves in *jaw-D,* likely due to petiole length differences only ($p < 0.05$, Wilcox test).
(D) Median length of the lamina measured from the petiole insertion point to the distal tip of the leaf is not significantly different between fully grown WT and *jaw-D* leaves ($p > 0.05$, Wilcox test).
(E) The median relative length between the first and second leaves within the same plant of WT and *jaw-D* is not significantly different, though *jaw-D* exhibits much more extreme values ($p > 0.05$ Wilcox test).
(F-G) The logarithm of leaf length (F) and leaf area (G) with plotting as in (A) demonstrate these values are increasing exponentially over the live imaging experiment.
Associated with Fig 1.

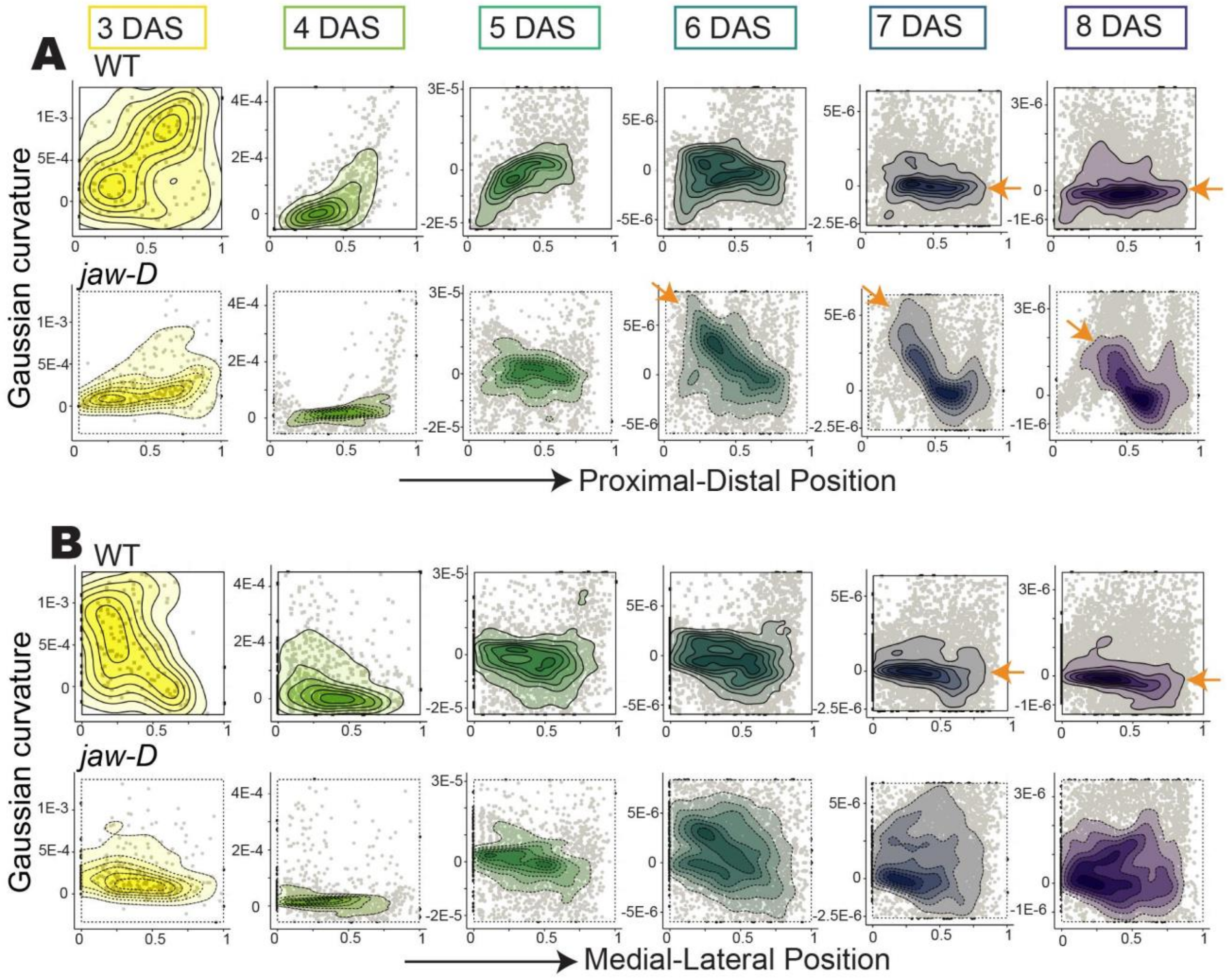


**Supplemental Fig. S8** ***jaw-D*** **leaves have high Gaussian curvature while wild type leaves flatten toward 0 Gaussian curvature**

(A-B) Quantification of the distribution of curvature along the proximal-distal (A) and medial-lateral (B) axes for all three replicates of WT (top, squares, solid lines) and *jaw-D* laminas (bottom, circles, dashed lines). A proximal-distal gradient is evident starting at 6 DAS in *jaw-D* leaves (orange arrow), whereas WT flattens starting at 7 DAS (orange arrows). In the medial-lateral direction, wild type flattens starting at 7 DAS (orange arrows), whereas jaw-D has broad lateral complexity of curvature. Note, as all leaves flatten, the Gaussian curvature changes dynamically over multiple orders of magnitude during the experiment, so the y-axis changes over time. Further, the petiole and margin are sources of extreme outliers in curvature. Therefore, the scale for the heatmaps and graphs representing each time point has been calculated to encompass the 85% range of curvature values around the mean for that time point. Associated with Fig 2.

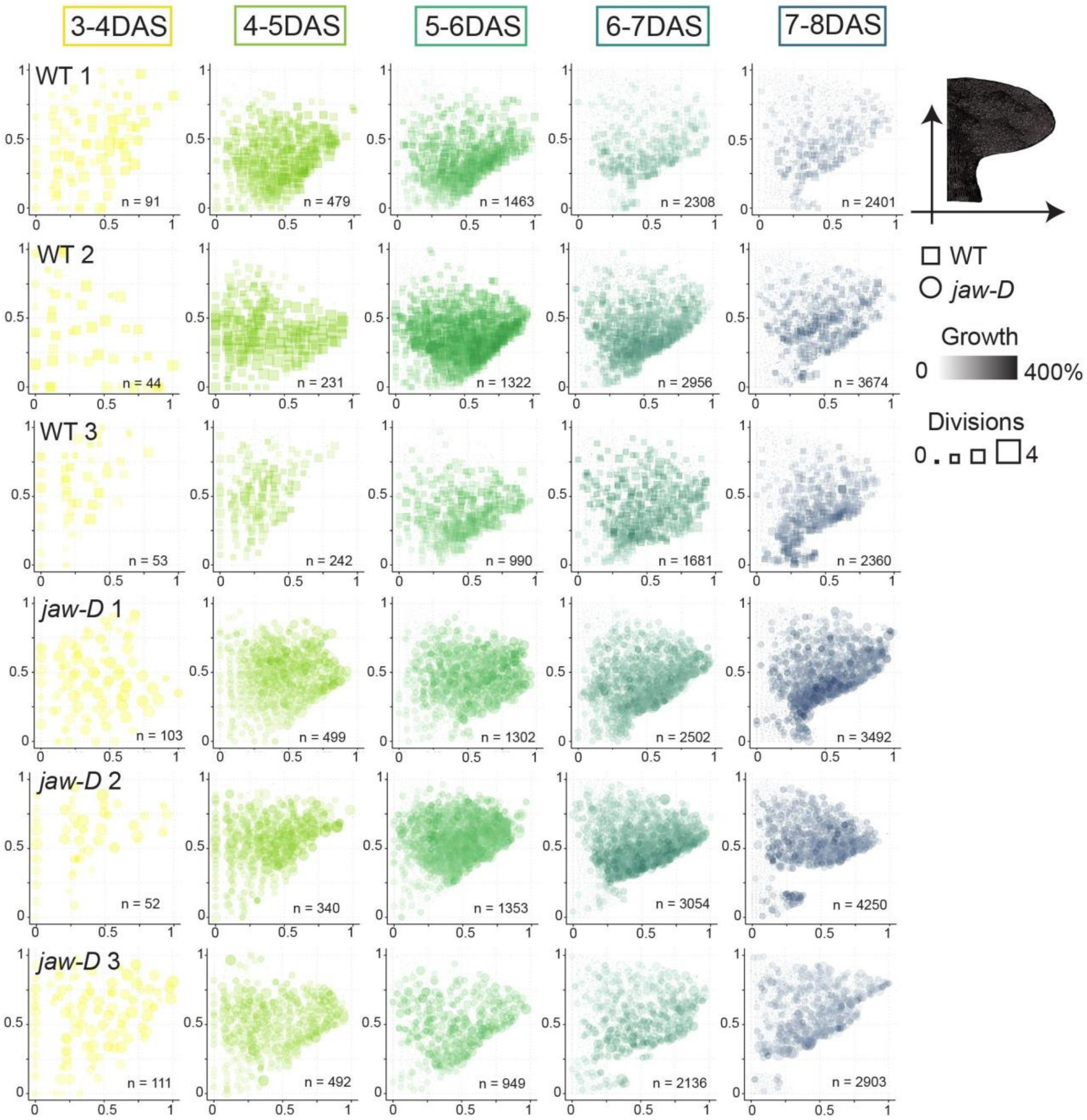


**Supplemental Fig. S9 Growth and division patterns are consistent between replicates**
Distribution of cell growth and division of each cell for one day intervals per whole leaf sample of WT (first three rows) and *jaw-D* (last three rows). n indicates the number of cells in each panel. The x and y axes represent medial-lateral and proximal-distal positions, respectively, normalized between 0 and 1 for the given leaf and time point. Colors indicate 24 hour time points. The size and color intensity of each point represents the number of divisions and areal growth, respectively, a cell has experienced between imaging days. Panels in S3 Fig. do not separate cells between the petiole and the lamina so the proximal-distal normalization is from the base of the petiole. Note, in the main text Fig. 3 the lamina lengths (without petiole) were calculated separately by selecting different ‘base cells’ to measure the distance from in

MorphoGraphX. Normalization was performed on these separated measures. Associated with Fig 3.

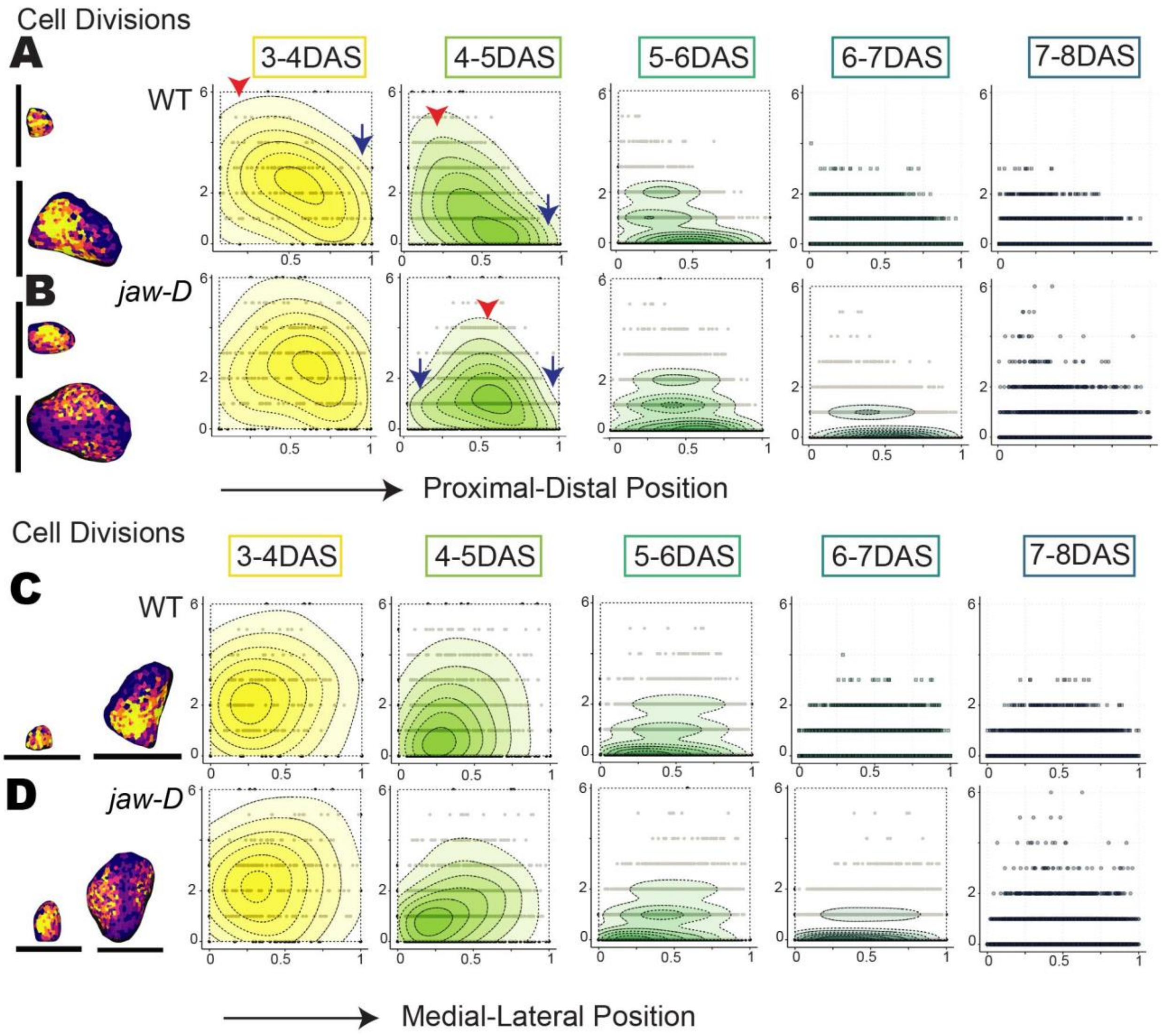


**Supplemental Fig. S10 Cell division patterns in WT and *jaw-D***

(A-B) Cell divisions over 24 hour intervals versus position along the proximal-distal axis for each cell in all three WT (A, solid lines, squares) lamina replicates or *jaw-D* (B, dotted lines, circles) replicates. Cell divisions are concentrated more distally in *jaw-D* than WT.

(C-D) Cell divisions over 24 hour intervals versus position along the medial-lateral axis for each cell in all three WT (C, solid lines, squares) lamina replicates or *jaw-D* (D, dotted lines, circles) Cell divisions are distributed more similarly between WT and *jaw-D* along the medial-lateral axis.

Use of density plots, coloring, axis selection and normalization as in Fig 4. Note, density plots in later time points were not calculated if the majority of cells were not dividing. In this case, only points representing individual cells are plotted.

Associated with Fig. 4.

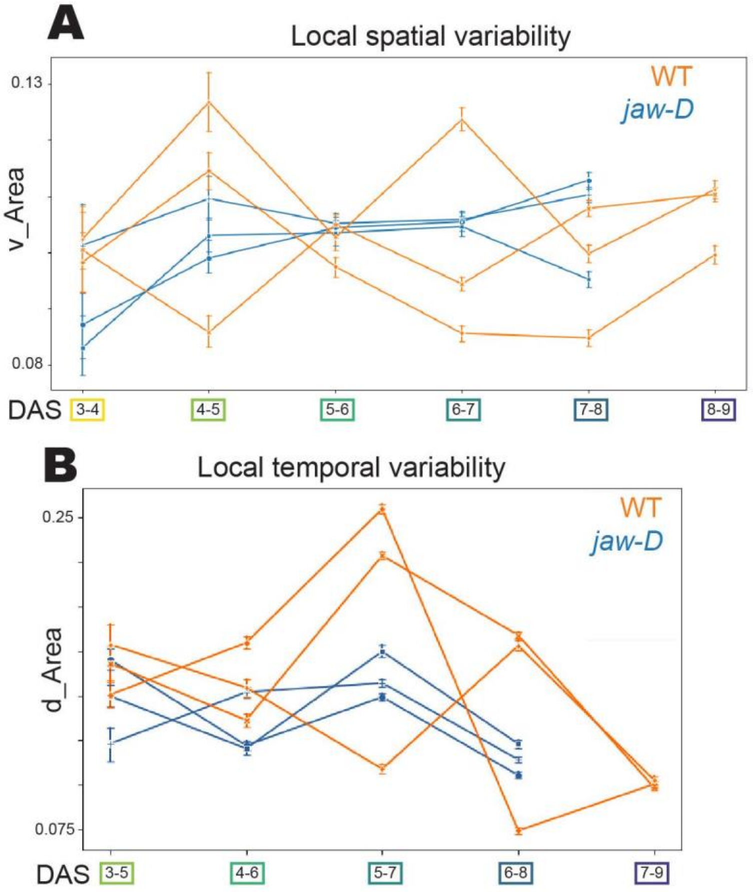


**Supplemental Fig. S11 Spatio-temporal analysis with stomata excluded**
(A-B) Quantification of the mean (A) local growth spatial variability or (B) local growth temporal variability for time windows as in main Figure 5C-D, with stomata excluded. Excluding stomata does not change patterns of growth heterogeneity observed between WT and *jaw-D.* WT and *jaw-D* local spatial and temporal growth variability are largely indistinguishable, though *jaw-D* replicates are generally tighter around the mean.

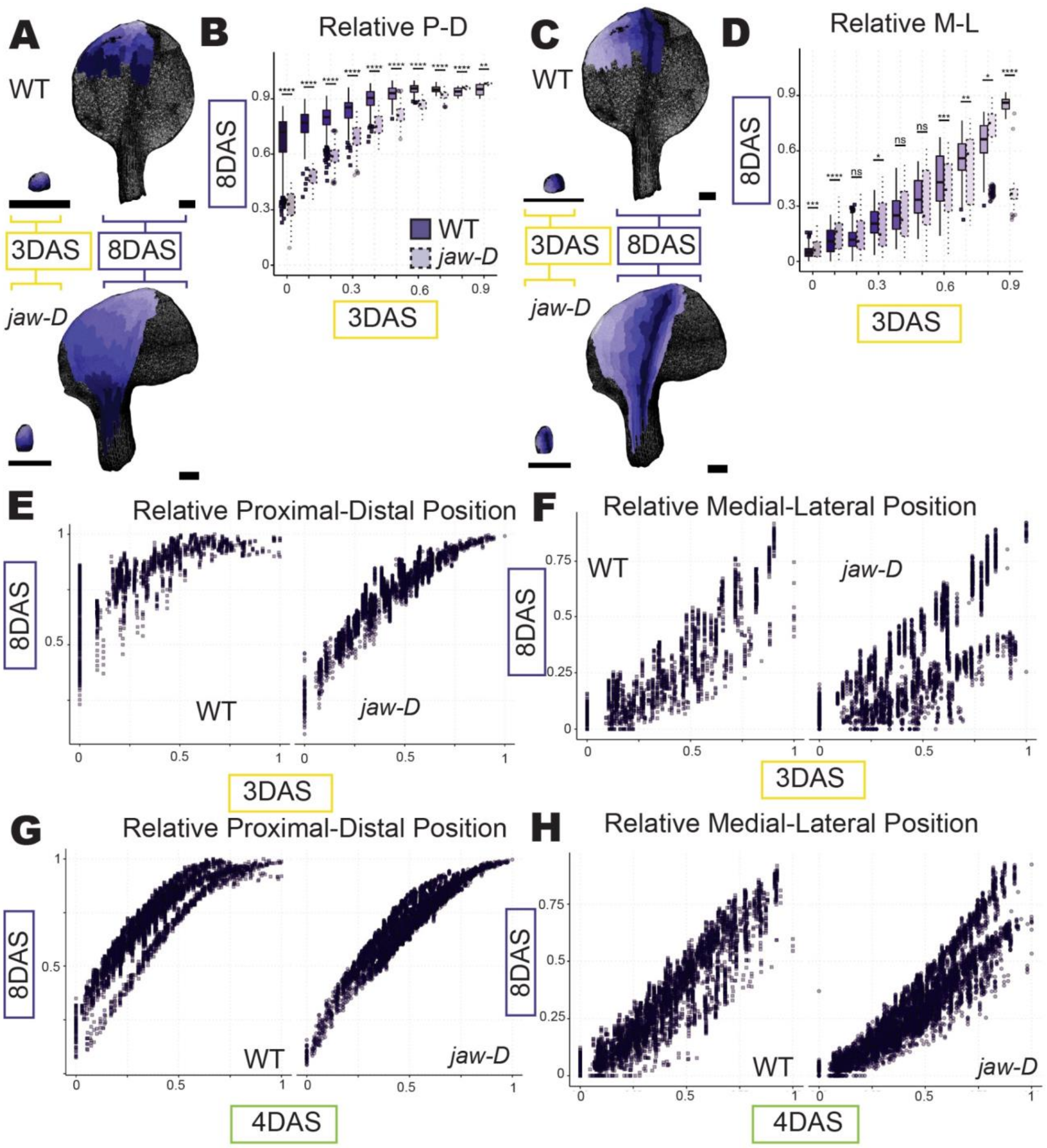


**Supplemental Fig. S12 Additional clonal analysis**

(A-D) Clonal analysis of WT (top) and *jaw-D* (bottom) clonal sectors mapped from 3 DAS to 8 DAS on sample leaf meshes in shades of purple along the proximal-distal (A) or medial-lateral (C) axis. Clonal analysis is as in Fig. 5, except that it starts at 3 DAS instead of 4DAS. (B,D) Quantification for all three whole leaf replicates. WT (left, dark shading, solid lines) and *jaw-D* (right, light shading, dotted lines). Boxes indicate the 75% interquartile range, middle bar represents median, whiskers represent outliers and squares (WT) or circles (*jaw-D)* represent extreme values. Nearly all 3 DAS proximal-distal bins have similar to enhanced results as 4 DAS with significant differences for 8 DAS locations between WT and *jaw-D,* while medial-

lateral contributions are more mixed (**** = p<.0001, ** = p <.01, * = p <.05, paired Student's t-test and Bonferroni correction). Final leaf contributions from WT seem to emerge mostly mid-leaf and a consistent stronger contribution from *jaw-D* lateral domains. Black scale bars = 200 µm. (E-H) Plots of all cells at 8 DAS versus their relative proximal-distal (E,G) or medial-lateral (F,H) position in the 3DAS (E-F) or 4DAS (G-H) leaf. Bifurcations in *jaw-D* medial-lateral positions could indicate tissue growth conflicts that induce rippling (F,H).
Associated with Fig. 6.

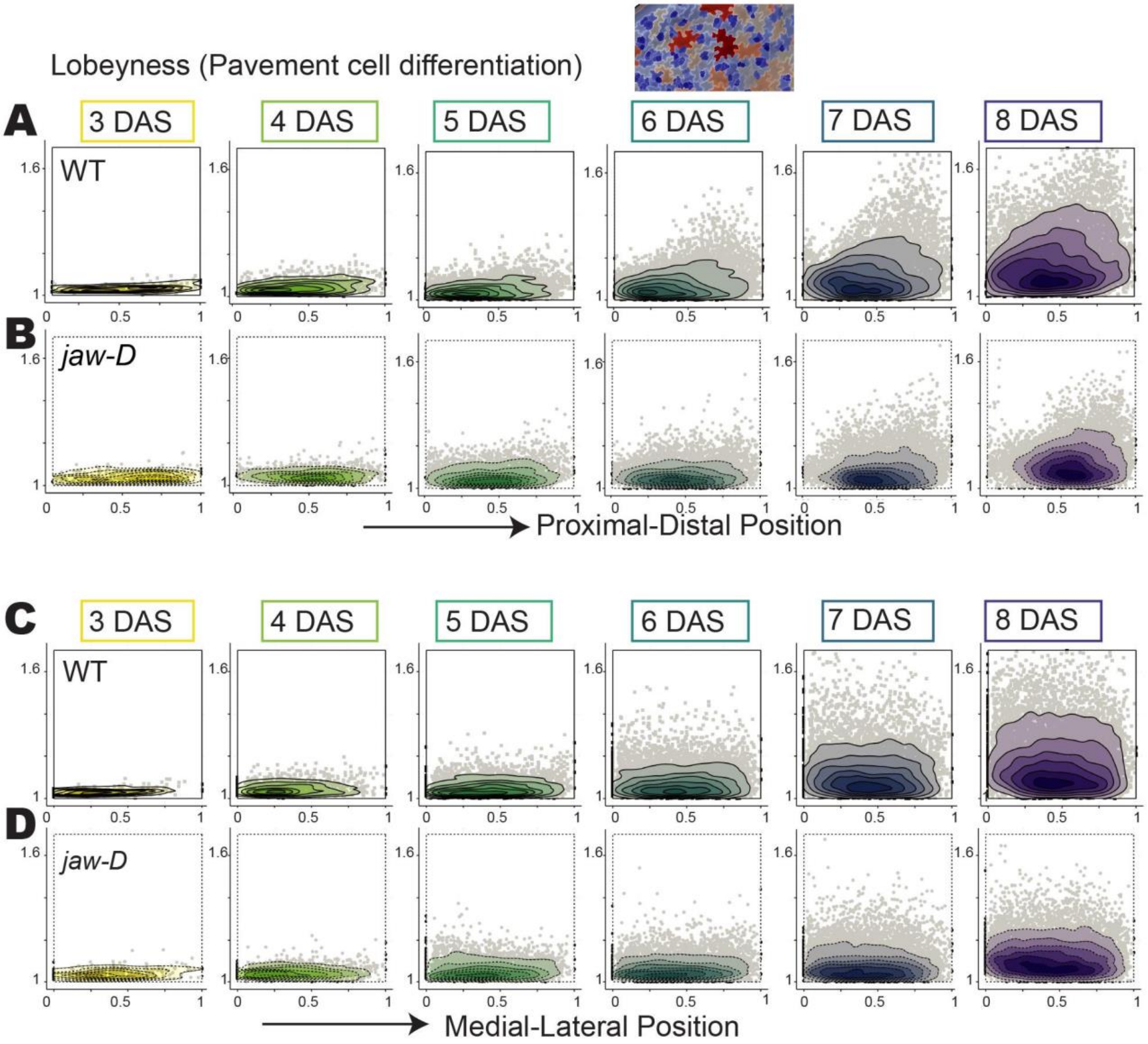


**Supplemental Fig. S13 Quantification of differentiation as measured by lobeyness**

(A-B) Lobeyness (the perimeter of a cell divided by its convex hull, a measure of pavement cell identity) versus position along the proximal-distal axis for each cell in all three WT (A, squares, solid lines) or *jaw-D* (B, circles, dotted lines) lamina replicates. Lobeyness is initiated at the distal tip of WT cells at 5 DAS and increases while propagating down to the leaf base. Increases in lobeyness are subtler, later and more disbursed in *jaw-D* samples.

(C-D) Lobeyness is more evenly distributed across the medial-lateral axis in WT(C) and *jaw-D* (D) with greater absolute magnitude in WT.

Use of density plots, coloring, axis selection and normalization as in Fig 4.

Associated with Fig. 7.

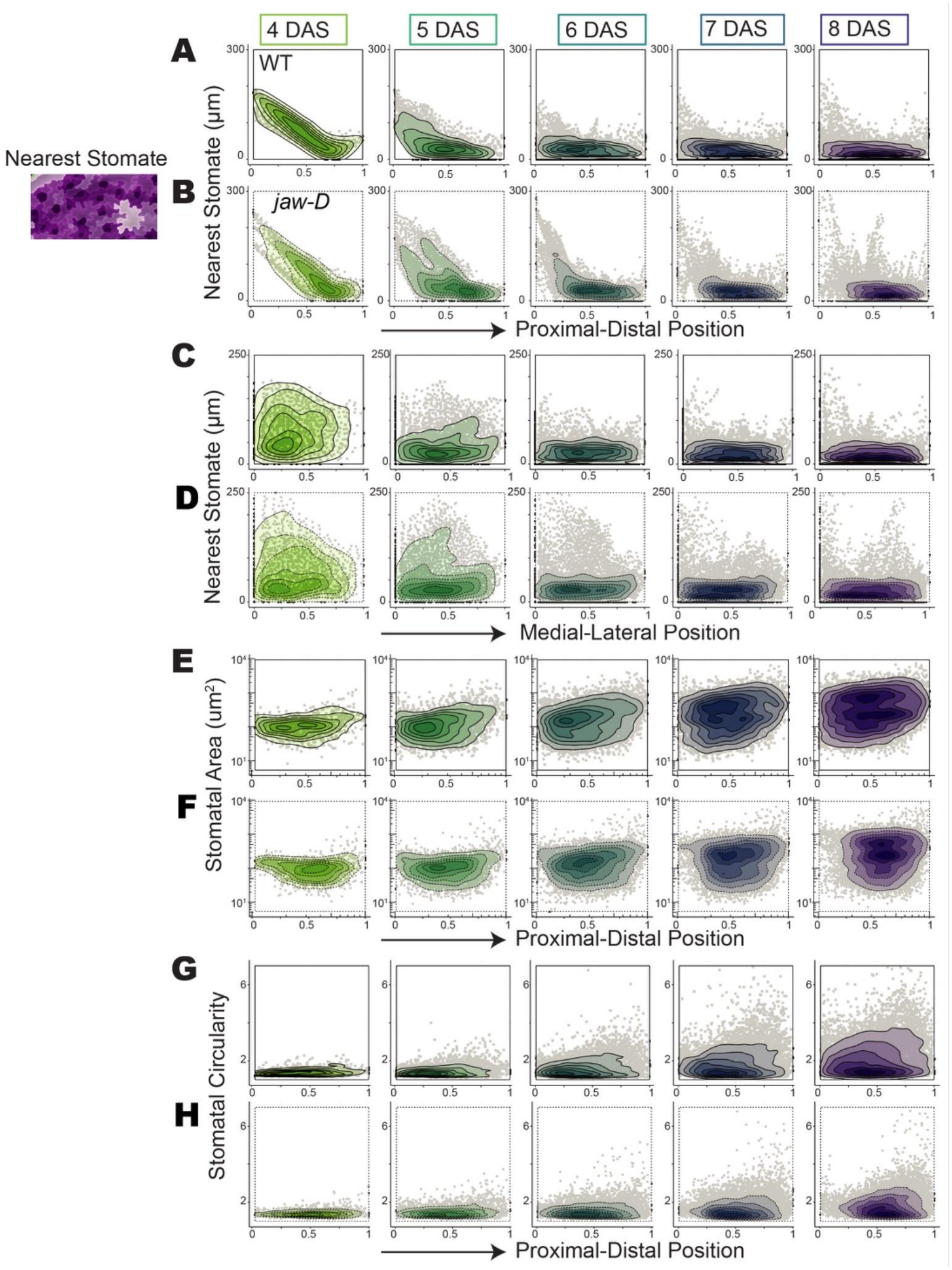


**Supplemental Fig. S14 Quantification of differentiation of stomata**

(A-D) Quantification of stomatal spacing (the distance in µm from pavement cells to the nearest stomata, the inverse of stomatal density) versus position along the proximal-distal axis (A-B) or the medial-lateral axis (C-D) and WT (A,C) and *jaw-D* (B, D). Note, stomata first initiate at 4DAS. Stomatal spacing is largely the same between WT and *jaw-D* along both axes. Stomatal spacing quickly tapers as stomata are initiated and become evenly distributed through the lamina.
(E-H) Area (E-F) or circularity (G-H) (perimeter/4pi*r^2) versus position along the proximal-distal axis for each stomate in all three WT (E,G, squares, solid lines) or *jaw-D* (F,H, circles, dotted lines) lamina replicates. (E) Stomatal area shows a distal maxima with a graded decrease towards the base for WT samples starting at 5 DAS. (F) Stomata in *jaw-D* have more even sizes. Note, as the stomatal areas cover many orders of magnitude over time, a log scale is used. (G) Stomatal circularity is further from 1 in WT in a graded pattern signifying anisotropic expansion from tip to base. (F) *jaw-D* stomata are rounder throughout the lamina. These differences demonstrate how stomata may be subject to tissue-level anisotropy and maturation processes only in WT while baseline initiation and differentiation for their physiological role is maintained in WT and *jaw-D*. Use of density plots, coloring, axis selection and normalization as in Fig 4.
Associated with Fig. 7.

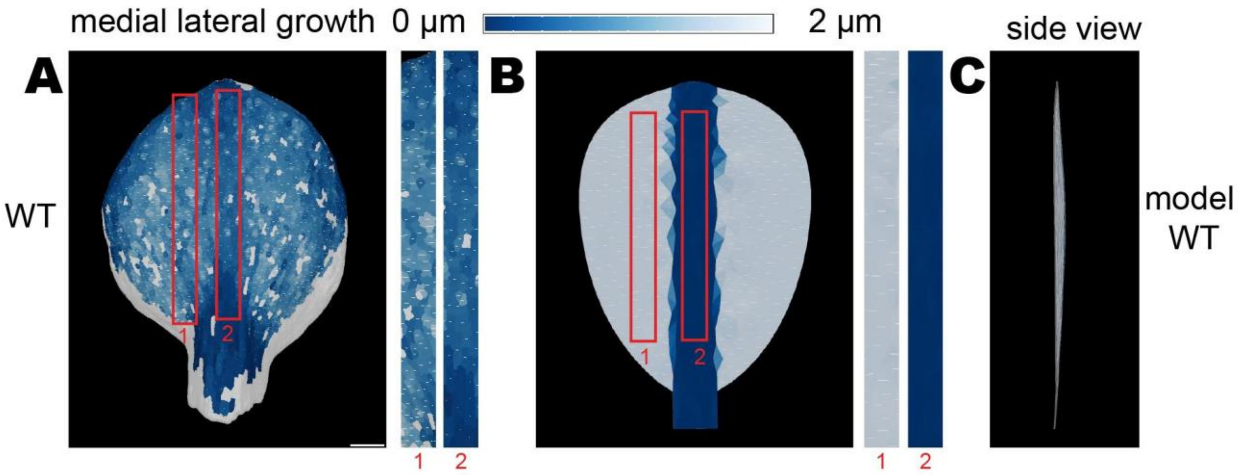


**Supplemental Fig. S15 The basipetal growth gradient does not cause a loss of flatness**

(A) Heat map of cell growth along the medial-lateral axis of the WT leaf from 3-4 DAS. High medial-lateral extension is colored white (2 µm) and no extension is colored blue (0 µm). The magnitude of growth along this medial-lateral axis is also represented by the length of the white line drawn in each cell. Two regions are magnified to make the cellular growth rates visible across the axis of the leaf. Note that the medial-lateral growth rates at each proximal-distal region of the leaf are similar, although the tip of the leaf is growing slower due to the basipetal growth gradient.
(B-C) Finite element model simulation of a leaf growth in which the medial-lateral growth is displayed. In the model, the midrib grows very little in width in comparison to the blade, however, this does not create a growth conflict because the whole length of the midrib does not grow. (C) A 90 degree rotation of the same leaf simulation (side view), showing that the leaf remains relatively flat despite small growth conflicts caused by the basipetal gradient. Associated with Fig. 8.

**S1 Video. Clonal lineage of 4 DAS and 8 DAS WT meshes in MGX**
Mesh representations of 4 DAS (right) and 8 DAS (left) meshes of sample WT leaf in MorphoGraphX software. Video begins showing 4 DAS sample with segmented cells indicated with different colored labels. Zoom out shows relative sizes and morphology changes from 4 DAS to 8 DAS. Parent labels are then displayed on 8 DAS mesh to indicate how cells labeled in 4 DAS mesh contributed to 8 DAS sample. Relatively simple, isotropic cells in an amorphous nub have become lobed pavement cells and stomata while the gross leaf morphology has changed to a distinct petiole and flattened, round blade. Black scale bar = 200 um.

**S2 Video. Clonal lineage of 4 DAS and 8 DAS *jaw-D* meshes in MGX**
Representations as in Video S1 for *jaw-D* 4 DAS (left) and 8 DAS (right) sample meshes. 8 DAS mesh in *jaw-D* exhibits enhanced proximal curvature, and smaller, less lobed cells. Clonal patches are also smaller. Black scale bar = 200 um.

**S3 Video. 4-8 DAS clonal lineages with animated growth in WT**
Clonal lineages traced from 4 DAS to 8 DAS on example WT mesh. Individual lineages indicated as different label colors. Unlabeled cells could not be traced between all time points. Proximal lineages tend to divide and expand many times and often anisotropically, while distal lineages are more isotropic, expand and divide less. Black scale bar = 200 um

**S4 Video. 4-8 DAS clonal lineages with animated growth in *jaw-D***
Clonal lineages for a *jaw-D* sample with representation as in Video S3. All lineages are generally more isotropic in expansion and divide less.

**S5 Video. Curvature cross-sections of WT and *jaw-D***
2.5D mesh representations of WT (left, top) and *jaw-D* (right, bottom) 8 DAS samples arranged in MorphoGraphX software. Meshes overlaid to compare curvature at comparable cross-section locations. Clipping plane (white lines) used to scroll through mesh sections to show proximal curvature increase in *jaw-D*. White scale bar = 200 um

**S6 Video. Simulation of WT leaf with uniform growth across the width of the leaf**
FEM simulations of leaf mesh growth as in Fig 8E-F, representing WT. Heatmap and white lines indicate proximal-distal growth (white high growth and blue no growth). The basipetal growth gradient has been imposed, but growth across the width is uniform, such that no growth conflicts are created and the leaf remains flat.

**S7 Video. Simulation of *jaw-D* leaf with symmetric faster growth in the blade than the midrib.**
FEM simulations of leaf mesh growth as in Fig 8G-H, representing *jaw-D*. Heatmap and white lines indicate proximal-distal growth (white high growth and blue no growth). The basipetal growth gradient has been imposed, and proximal-distal growth of the midrib is lower than growth in the blade (symmetric on both sides), which causes growth conflicts and curvature of the leaf.

**S8 Video. Simulation of *jaw-D* leaf with asymmetric faster growth in the blade than the midrib.**

FEM simulations of leaf mesh growth as in Fig 8I-J, representing *jaw-D*. Heatmap and white lines indicate proximal-distal growth (white high growth and blue no growth). The basipetal growth gradient has been imposed, and growth of the midrib is lower than growth in the blade (with different amounts of growth on each side causing asymmetry), which causes growth conflicts and curvature of the leaf.

**S9 Video. Simulation of WT leaf with basipetal medial lateral growth**

FEM simulations of leaf mesh growth as in Fig 9B-C, representing WT. Heatmap and white lines indicate medial-lateral growth (white high growth and blue no growth). The basipetal growth gradient has been imposed as well as very low medial-lateral growth of the midrib. Although the basipetal growth gradient causes a small amount of growth conflict in the medial-lateral orientation, the anisotropy of the growth minimizes the curvature generated and the leaf remains relatively flat.
