## Supporting Information for "Cell growth rates coordinate across the width of the leaf to remain flat"

To accompany

#### 1 One-dimensional model of leaf length

In this section, we develop a simple one-dimensional model to capture the length of the leaf over time. Our goal is to explore how contrasting spatial growth profiles for WT and jaw-D genotypes can lead to overall similar leaf length dynamics. These differences are clearly seen in Fig. 3A-B on days 4-6, where cell growth shows a sharp decline with the proximal-distal location in WT. In contrast, the spatial growth profiles are relatively flat with respect to the spatial position in jaw-D.

We formulate a discrete-time model where time $t = 0, 1, 2, \ldots$ corresponds to days and $L_t$ is the leaf length at $t$ days after sowing. We assume that cells are arranged linearly in a one-dimensional lattice from position $i = 1, 2, \ldots L_t$. Let $p_{i,t}$ denote the probability of division for a cell at location $i$ between time $t$ and $t+1$. Then the leaf length at time $t+1$ is given by

$$L_{t+1} = \text{Round}\left[\sum_{i=1}^{L_t} 1 + p_{i,t}\right]$$

where Round is the nearest-integer operation. In this model, the probability $p_{i,t}$ is analogous to the cellular growth rate. Starting from a single cell at position $i = 1$ at time $t = 0$, the length grows exponentially in the beginning. This is captured by having $p_{i,t} = 1$ as long as the leaf length $L_t$ is below a threshold $\bar{L}$. As the leaf dynamics exits this initial exponential phase, $p_{i,t}$ becomes a function of both time and spatial location, and also differs between WT and jaw-D.

For WT we consider the following functional form:

$$p_{i,t} = \frac{p_{0,t}}{1 + (ki)^2}$$

that decreases with increasing distance from the leaf base $i$, where $k$ is a positive constant. Here $p_{0,t}$ is the growth at the leaf base which decreases over time, and this decrease is essential to eventually stop all growth as the leaf length approaches its steady-state value $L_\infty$. We consider the following form for the growth rate at the leaf base that decreases with increasing length $L_t$:

$$p_{0,t} = \frac{\left(1 - \frac{L_t}{L_\infty}\right)^2}{\left(1 - \frac{\bar{L}}{L_\infty}\right)^2}$$

Given the flat growth profile seen in jaw-D, the probability of cell division is assumed to be independent of $i$ and given by

$$p_{i,t} = \frac{\left(1 - \frac{L_t}{L_\infty}\right)^2}{2\left(1 - \frac{\bar{L}}{L_\infty}\right)^2}$$

The model simulations are shown in Fig. 4C-D with $\bar{L} = 50a.u.$, $L_\infty = 500a.u$, $k = 0.01$, and show similar length dynamics between the two genotypes.

### 2 Cellular Fourier transform

Here we define the discrete Laplace operator, we explain how we built the Fourier harmonics based on this Laplace operator, we define cell growth and deformation, and detail how we performed the spectral analysis based on CFTs. The theoretical basis of the CFT may be found in [4].

#### 2.1 Discrete Laplace operator

To compute the Laplace operator, we consider the cells' minimal surfaces whose contours are the cells' contours. For each cell, we project its contour on a plane that is perpendicular to its surface vector. The contour being polygonal, the surface vector can be written $1/2 \sum_n \vec{r}_n \wedge \vec{r}_{n+1}$ where the sum is over the contour vertexes, $\vec{r}_n$ is their position, $n$ indexes the position around the contour and $\wedge$ is the exterior product. We then triangulate the surface enclosed in the projected contour using the MESH2D Matlab package [2, 1]. To obtain a mesh approximating the 3D minimal surface, we then interpolated positions in the mesh along the surface vector using cell contours. The area $S_{i,t}$ for cell $i$ at time $t$ is then computed as the sum of areas of triangles in the triangulation, $S_{i,t} = \sum_m^{(i,t)} dS_m$, where $m$ spans triangles of cell $i$ at time $t$ and $dS_m$ is the area of triangle $\#m$. The tissue is made of $N$ cells that are followed from $t$ to $t+1$.

The discrete Laplace operator is a square matrix of size $N \times N$ and its components are given by

$$\bar{L}_{ij,t} = \delta_{ij} - \bar{W}_{ij,t},$$
$$\text{where } \bar{W}_{ij,t} = \sqrt{\frac{S_{i,t}}{S_{j,t}}} \frac{\sum_m^{(i,t)} dS_m \sum_n^{(j,t)} dS_n \exp(-d_{mn}/(5\,\ell_c))}{\sum_m^{(i,t)} dS_m \sum_j \sum_n^{(j,t)} dS_n \exp(-d_{mn}/(5\,\ell_c))}.$$

Here indices $i = 0, 1, ..N-1$ and $j = 0, 1, ..N-1$ span the $N$ cells of the tissue. $d_{mn}$ is the distance between triangle $m$ from cell $i$ and triangle $n$ from cell $j$, both considered at time step $t$. The unit of length is mean cell size $\ell_c = \sqrt{S_t/N}$, where $S_t$ is the surface of the tissue at time $t$ and $N$ is the number of cells. Here we took the width $5l_c$ for the coarse Laplace operator.

#### 2.2 Fourier harmonics

We start from the singular value decomposition of the Laplace operator $\bar{L}$, which yields left singular vectors $V$, right singular vectors $U$, and the singular values $\hat{L}_k$:

$$\bar{L}_{ij,t} = \sum_{k=0}^{N-1} \hat{L}_k \, V_{ki} U_{kj}.$$

The value taken by the $k^{\text{th}}$-harmonic in cell $i$ at time step $t$ is $1/S_{i,t} U_{ki}$. The spatial frequency of the $k^{\text{th}}$ harmonic is given by $f_k = 1/(5\pi)\, Q(\hat{L}_k)$, with $Q(l) = \sqrt{(1-l)^{-2/3} - 1}$. The harmonics are indexed so that their index grows with the wave number.

#### 2.3 Cell deformations & growth

To define the deformation of a cell $i$, we look for the linear transformation which best deforms the cell's contour at time $t$ into it's contour at time $t+1$ (the contour of daughter cells if the cell $i$ divides from $t$ to $t+1$). Both cell contours are first projected perpendicularly to their surface vectors. The coordinates $\vec{r}_{n,t}$ and $\vec{r}_{n,t+1}$ of their vertices are considered in two orthogonal basis of their projection planes, centered at the cell center. The best deformation is obtained in finding the matrix $M_i$ which minimizes a cost function $d(M_{i,t} \cdot \vec{r}_{n,t}, \vec{r}_{n,t+1})$. The cost function $d$ is built from the the Euclidean distance $|.|$:

$$d(\vec{r}_{n,t}, \vec{r}_{n,t+1}) = \sum_{j}^{(i)} \sum_{n \in \mathcal{V}_{j,t}} \text{Min}_{n' \in \tilde{\mathcal{V}}_{j,t+1}} |\vec{r}_{n,t} - \vec{r}_{n',t+1}|^2 + \sum_{j}^{(i)} \sum_{n' \in \tilde{\mathcal{V}}_{j,t+1}} \text{Min}_{n \in \mathcal{V}_{j,t}} |\vec{r}_{n,t} - \vec{r}_{n',t+1}|^2,$$

where the sum $\sum_j^{(i)}$ is over the neighboring cells, $\mathcal{V}_{j,t}$ are the vertex of the cell's contour also neighbor with $j$ at $t$, and $\tilde{\mathcal{V}}_{j,t+1}$ the vertex to the corresponding segment of the cell contour at time $t+1$. A polar decomposition can be made of the matrix $M_{i,t} = R_{i,t} \cdot D_{i,t}$, where $R$ is an arbitrary rotation and $D_{i,t}$ gives the cell deformation in the reference frame of the cell contour at time $t$. If this matrix describes well the cell's deformation, its eigenvectors give the principal and secondary direction of growth, the relative difference between the first and the second eigenvalues $d_{i,t}^{(1)}$ and $d_{i,t}^{(2)}$ gives an estimate for the cell growth anisotropy, while their product should be $1 + G_{i,t}\,\Delta t$ with the $G_{i,t}$ areal growth rate and $\Delta t$, the delay between two time steps. The cell's areal growth rate $G_{i,t} = (\tilde{S}_{i,t+1}/S_{i,t} - 1)/\Delta t$ can also be calculated from its surface area $S_{i,t}$ at time $t$, and $\tilde{S}_{i,t+1}$ at time $t+1$ ( if the cell $i$ divides from $t$ to $t+1$, $\tilde{S}_{i,t+1}$ is the surface area of the daughter cells). We found good agreement between this last expression of $G_{i,t}$ and $d^{(1)}_{i,t} d^{(2)}_{i,t}$. The matrices $D_{i,t}$ being defined in bases of each cells' tangent planes, we then consider $D^{(3D)}_{i,t}$ the deformation matrices in a unique basis of the 3D space, the deformation along the surface vector being taken equal to zero.

##### 2.4 Spectral analysis

We applied our spectral analysis to the areal growth rate and the deformation. For the areal growth rate we used the formula $G_{i,t} = (\tilde{S}_{i,t+1}/S_{i,t} - 1)/\Delta t$. The $k^{\text{th}}$ CFT coefficient for the areal growth rate is then $\hat{G}_{k,t} = \sum_i U_{ki} G_{i,t} \sqrt{S_{i,t}/S_t}$ where $S_t$ is the total area $S_t = \sum_i S_{i,t}$. Here we use a convention that differs from [4] by a multiplicative factor $1/\sqrt{S_t}$ in the definition of the CFT. This makes the interpretation of CFTs simpler: they have the same dimensions (units) as the original signal (here growth) and the first coefficient is equal to the average signal. For the deformation matrix in 3D $D^{(3D)}_{i,t}$, we computed the CFT, $\hat{D}_{k,t} = \sum_i U_{ki} D^{(3D)}_{i,t} \sqrt{S_{i,t}/S_t}$.

Spectra are obtained in representing CFTs as function of the space frequency of the harmonics they are associated to. In the case of deformation, representing the spectra is complicated by the high number of the matrix's components, and another difficulty is to compare the spectra of different tissues whose orientations are arbitrary. To analyze the deviation of growth from isotropy we can nevertheless consider the difference $\hat{d}^{(1)}_{k,t} - \hat{d}^{(2)}_{k,t}$ between the two higher eigenvalues of $\hat{D}_{k,t}$, $\hat{d}^{(1)}_{k,t}$ and $\hat{d}^{(2)}_{k,t}$ which we represent once again as function of the spatial frequency of the harmonics they are associated to. To compare different tissues and genotypes, and since Fourier harmonics are specific to each tissue, we smooth spectra to obtain an estimate of spectral densities. The smoothing is made for every tissue with a Gaussian kernel of width 0.16 and is relative to the spatial frequency $f_k$. We also merge the spectra of identical stage and genotype. We estimate the standard error assuming the CFTs to be statistically independent and to be well described by distributions varying smoothly with respect to the spatial frequency.

### 3 Finite element model simulations

Leaves were composed of linear wedge elements with 9 Gaussian integration points [8]. The material law was described by an isotropic Saint Venant-Kirchoff material, with Young modulus $E = 100MPa$ and Poisson's ratio $\nu = 0.3$.

The Green-Lagrange strain tensor was used as measure of the deformation and the system equilibrium was calculated iteratively via an implicit Euler integration minimizing the total strain energy of the leaf [7, 6]. The simulation was considered to be at equilibrium when the the sum of the squared residual forces and the maximal residual force were below assigned thresholds. To speed up the calculations of elemental reaction forces and the elemental second derivative, isoparametric coordinates were adopted to describe the wedge reference configuration as illustrated in [3].

Growth was implemented by increasing the size of the reference configuration of each wedge and specified only in the plane parallel to the leaf surface. No growth is defined in the orthogonal direction (the thickness of the wedge reference configuration remains constant). Thus, any out-of plane deformation of the leaf is an emergent property of the model.

In order to prescribe in-plane growth, starting from the reference configuration of each wedge element, a special *midplane triangle* was defined as the intersection between the wedge and a plane crossing its quadrilateral faces at their half-height (see Fig. SI1). $K_{\text{par}}$ and $K_{\text{per}}$ specify the amount of growth parallel and orthogonal to the vector **KPar** respectively, with both directions lying on the midplane triangle [5]. For this work, **KPar** was set parallel to the leaf's proximal-distal axis.

To compute the grown configuration of the reference wedge, the following steps are performed:

- Both reference wedge and midplane triangle are translated so that their first vertex lies in the origin.
- The midplane triangle is rotated in the x-y plane by means of the 3D rotation matrix $\mathbf{R_{3To2D}}$ which rotates the triangle norm $\mathbf{N}$, taken as the cross product of its first and second sides, into a vector parallel to the z-axis and positively oriented with respect to it. The same rotation matrix is applied also to the reference wedge coordinates.
- The direction **KPar** is stored with the reference configuration by means of the angle $\alpha$ it makes with respect to the first side of the midplane triangle (both wedge and triangle sides are consistently oriented), and the triangle is rotated in the 2D plane by means of the rotation matrix $\mathbf{R_{KParToY}}$ so that the vector **KPar** is aligned with the y-axis (see Fig.SI1).
- The 2D rotation matrix $\mathbf{R_{KParToY}}$ is extended into a 3D matrix in the following way:

$$\mathbf{R_{KParToY3D}} = \left(\begin{array}{cc|c} \multicolumn{2}{c}{\multirow{2}{*}{$\mathbf{R_{KParToY}}$}} & 0 \\ & & 0 \\ \hline 0 & 0 & 1 \end{array}\right)$$

  and is then applied to the reference wedge.
- Since the midplane triangle is rotated so that **KPar** is parallel to the y-axis, it is now possible to express the specified growth in the form of a 2D

diagonal tensor

$$\mathbf{F_g} = \begin{pmatrix} 1 + K_{\text{per}} dt & 0 \\ 0 & 1 + K_{\text{par}} dt \end{pmatrix}$$

and $\mathbf{F_g}$ is applied to the midplane triangle coordinates $\mathbf{m_1}, \mathbf{m_2}, \mathbf{m_3}$ and the new, updated angle formed by **KPar** with the first triangle side $\mathbf{m_2 m_1}$ is computed and stored with the reference configuration.

- $\mathbf{F_g}$ is also turned into a 3D tensor

$$\mathbf{F_G} = \left(\begin{array}{cc|c} \multicolumn{2}{c|}{\multirow{2}{*}{$\mathbf{F_g}$}} & 0 \\ & & 0 \\ \hline 0 & 0 & 1 \end{array}\right)$$

and is applied to the reference wedge coordinates.

- The grown reference wedge is translated again so that its first vertex is back at its initial position. This translation procedure reduces numerical problems when computing element strain energy.

After each growth step, the mechanical equilibrium was computed, and the leaf deformed into the new equilibrium configuration.

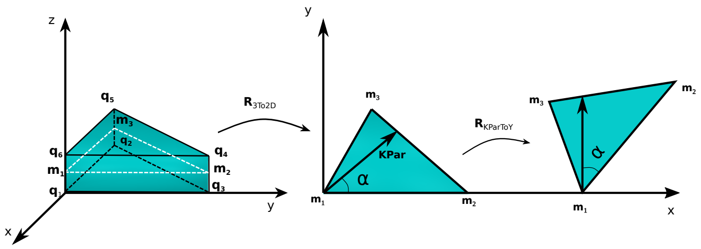


Figure SI1: Schematic drawing of how prescribed growth is assigned to the reference wedge element. **Left**: The wedge element is translated with its first node $q_1$ in the origin. The mid triangle with nodes $m_1$, $m_2$, $m_3$ is defined. **Right:** The midplane triangle is translated onto the x-y plane via the rotation matrix $\mathbf{R}_{3To2D}$ which also aligns its first side with the x-axis. The midplane triangle carries the information about the intrinsic main growth direction $K_{\text{par}}$ by means of the angle $\alpha$ it forms with the first triangle side $\mathbf{m}_1\mathbf{m}_2$. The midplane triangle is then rotated by the 2D matrix $\mathbf{R}_{KParToY}$ into the plane so that **KPar** is parallel to the y-axis.

### References

[1] Darren Engwirda. "Locally optimal Delaunay-refinement and optimisation-based mesh generation". PhD thesis. University of Sydney, 2014.

[2] Darren Engwirda. "Unstructured mesh methods for the Navier-Stokes equations". Undergraduate thesis. University of Sydney, 2005.

[3] Pavol Federl. *Modeling fracture formation on growing surfaces.* Citeseer, 2002.
