## Supplementary figures and images for "Cell growth rates coordinate across the width of the leaf to remain flat"

### Video S3

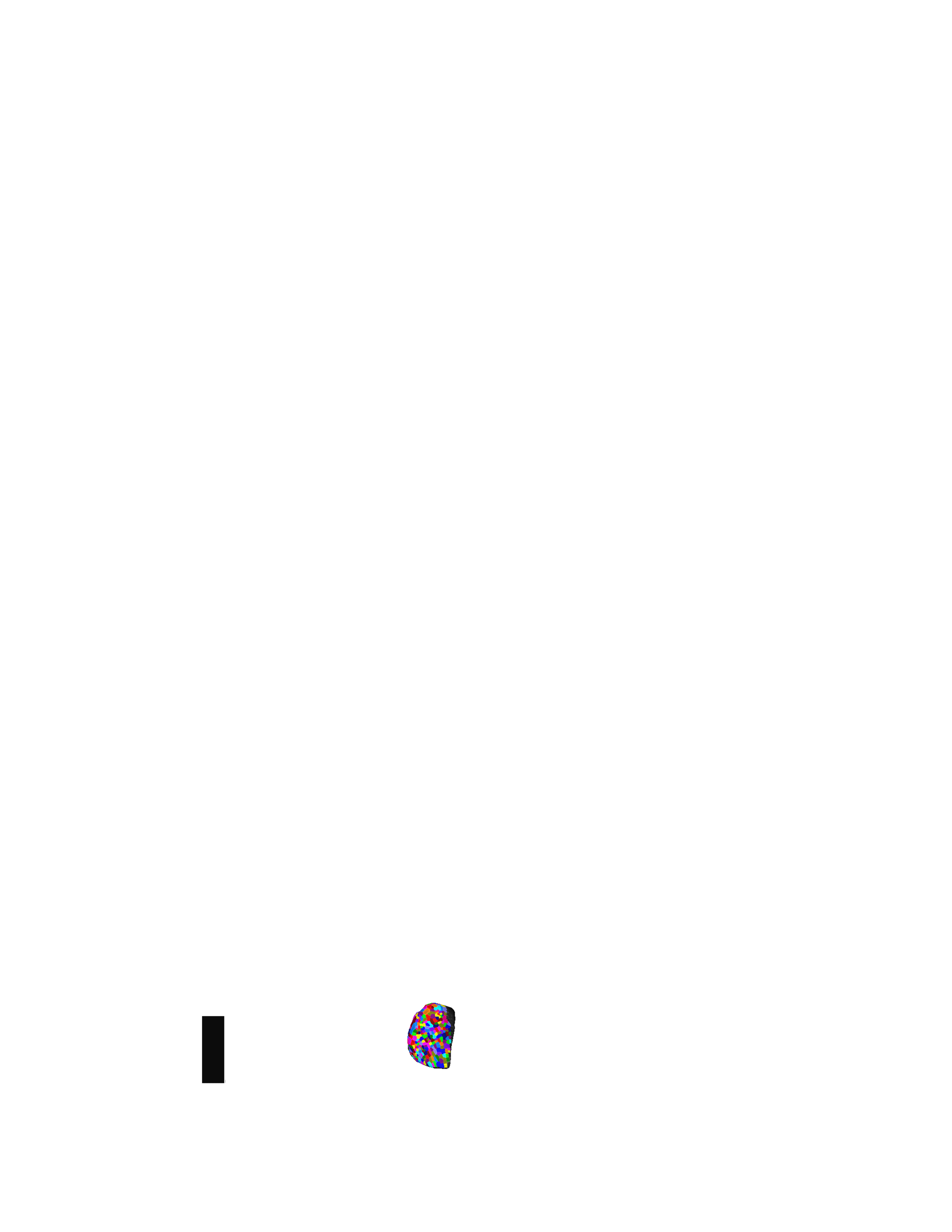

### Video S4

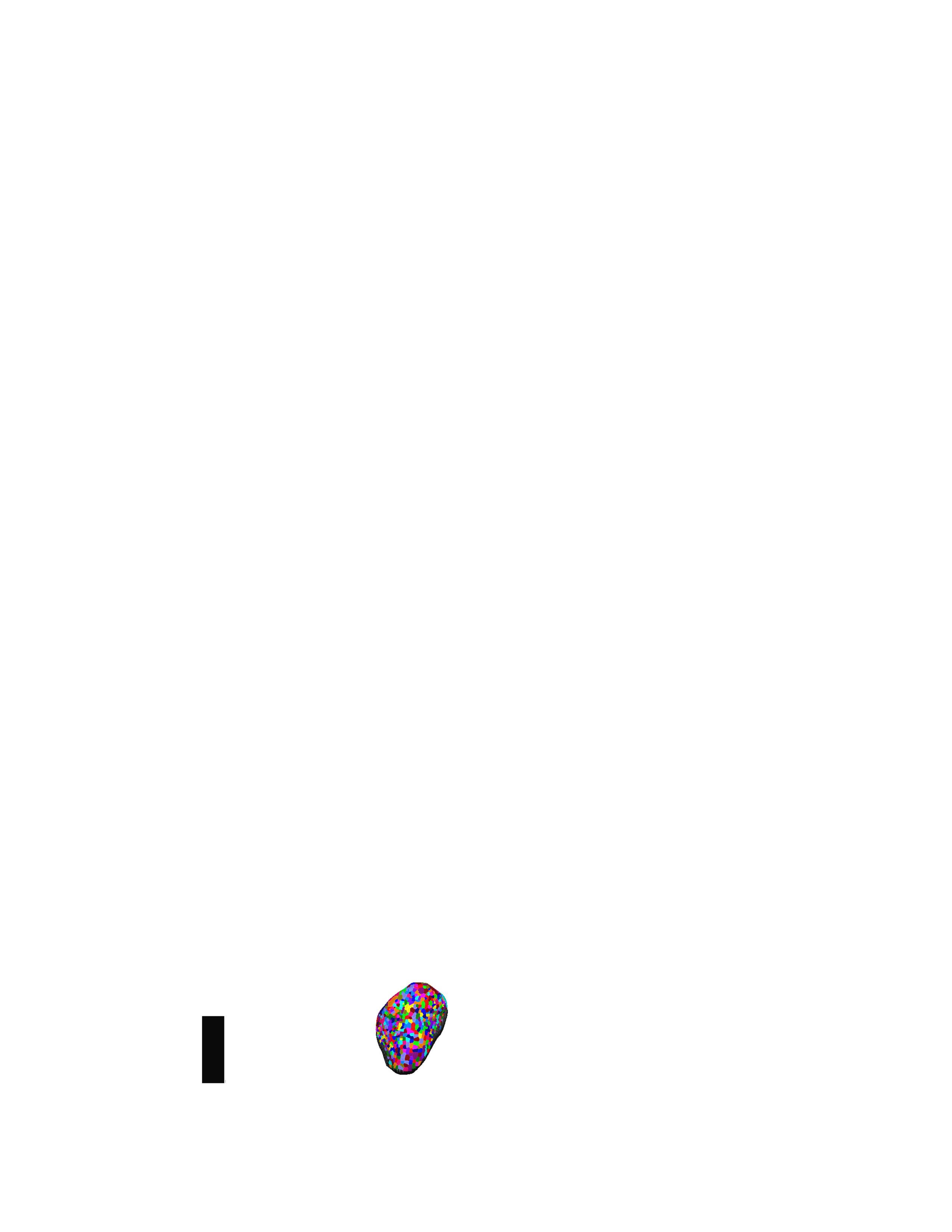
